# Endothelial cell invasiveness is controlled by myosin IIA-dependent inhibition of Arp2/3 activity

**DOI:** 10.1101/2020.09.08.287466

**Authors:** Ana M. Figueiredo, Pedro Barbacena, Ana Russo, Silvia Vaccaro, Daniela Ramalho, Andreia Pena, Aida P. Lima, Rita R. Ferreira, Fatima El-Marjou, Yulia Carvalho, Francisca F. Vasconcelos, Ana-Maria Lennon-Duménil, Danijela M. Vignevic, Claudio A. Franco

## Abstract

Sprouting angiogenesis is fundamental for development and contributes to multiple diseases, including cancer, diabetic retinopathy and cardiovascular diseases. Sprouting angiogenesis depends on the invasive properties of endothelial tip cells. However, there is very limited knowledge on the mechanisms that endothelial tip cells use to invade into tissues. Here, we prove that endothelial tip cells use long lamellipodia projections (LLPs) as the main cellular protrusion for invasion into non-vascular extracellular matrix. We show that LLPs and filopodia protrusions are balanced by myosin-IIA (MIIA) and actin-related protein 2/3 (Arp2/3) activity. Endothelial cell-autonomous ablation of MIIA promotes excessive LLPs formation in detriment of filopodia. Conversely, endothelial cell-autonomous ablation of Arp2/3 prevents LLPs development and leads to excessive filopodia formation. We further show that MIIA inhibits Rac1-dependent activation of Arp2/3, by regulating the maturation state of focal adhesions. Our discoveries establish the first comprehensive model of how endothelial tip cells regulate its protrusive activity and will pave the way towards new strategies to block invasive tip cells during sprouting angiogenesis.

## Introduction

The formation of new blood vessels from pre-existing ones – sprouting angiogenesis - is a fundamental process in health and disease (1, 2). Sprouting angiogenesis depends on the ability of a specialised endothelial cell - the tip cell – to invade and migrate into tissues (3), yet, we still lack basic understanding of how endothelial tip cells migrate and invade into tissues. During three-dimensional (3D) cell migration, endothelial tip cells, as other mesenchymal-like cells, such as fibroblasts and cancer cells, adopt a vast array of polarized membrane protrusions, which are essential for invasion and guidance (4–7). Membrane protrusions, such as lamellipodia and filopodia, are driven by actin dynamics (6, 8). Formin-dependent linear actin arrays promote filopodia formation, whilst actin-related protein 2/3 (Arp2/3) complex-dependent dendritic actin arrays promote lamellipodia formation (6). In addition, non-muscle myosin-II (MII)-dependent contractility was shown to inhibit protrusions and to promote membrane blebbing, through cortical tension (9, 10), whilst a recent study showed that MIIA, a specific MII isoform, could also promote filopodia stability (11). The balance between these different filamentous actin networks and their regulators shapes the types of cellular protrusions formed and hence the mode of migration. Vascular endothelial growth factor (VEGF) signaling is a master regulator of tip cell biology, and it was shown to regulate endothelial tip cell invasion and sprouting angiogenesis (1, 2, 12). CDC42 and RAC1, downstream of VEGF, are master regulators of endothelial tip cell membrane protrusions (13–15). VEGF also activates serum response factor (SRF), a transcription factor highly expressed by endothelial tip cells. Together with its co-factors myocardin-related transcription factor (MRTFs), SRF regulates endothelial invasive behavior by promoting filopodia formation (16–18). In addition, focal adhesions (FAs) are also crucial for EC migration and invasion, and deletion of integrin-linked kinase (ILK) or integrin β1 (ITGB1) leads to severe sprouting angiogenesis defects and changes in tip cell protrusive activity (19, 20). Despite their central role in tip cell invasion, it remains to be determined how actomyosin and its regulators affect cell shape and protrusive behaviour, and how different membrane protrusions influence endothelial cell migration *in vivo*. For instance, by using low doses of latrunculin B, a pharmacological inhibitor of actin polymerization, a recent report proposed that filopodia are dispensable for endothelial cell guidance and migration in zebrafish (21), generating controversy in the field.

Here, we investigated the specialized membrane protrusions on endothelial tip cells and their physiological relevance for endothelial cell invasion and angiogenesis in vivo, and generated an integrative model of how endothelial tip cells invade into tissues.

## Results

SRF/MRTFs activity was shown to regulate non-muscle myosin-II (MII) levels in endothelial cells (16). Endothelial cells express mostly two isoforms of MII, MIIA and MIIB, which are coded by Myh9 and Myh10, respectively (http://betsholtzlab.org/VascularSingleCells/database.html; (22)). To evaluate the function of MII in endothelial tip cells, we first investigated the location and expression levels of MIIA and MIIB. We first focused on MIIA. We found that endothelial tip cells express higher levels of MIIA than stalk cells, the cells juxtapose to tip cells that participate in the invading vascular sprout (Fig.1A,B, Sup.Fig.1A,B, and Sup.Videos 1 and 2). We further confirmed these observations by imaging fluorescently-tagged MIIA, using the Myh9-eGFP transgenic animals (Sup.Fig.2A) (23). In addition to higher expression, we found that MIIA was enriched at the base of filopodia protrusions (Fig.1A,C, and Sup.Fig.2B,C). MIIB followed a similar pattern to MIIA. It is highly expressed by endothelial tip cells and it is enriched at the base of filopodia (Sup.Fig.2C). This spatial distribution of MII isoforms correlates with high levels of phospho-myosin light chain (pMLC) and filamentous actin (Sup.Fig.2B-D). These results suggest that endothelial tip cells have high levels of actomyosin contractility at their leading edges, in particular at the base of filopodia. To evaluate the function of MIIA and MIIB in endothelial cells, we deleted Myh9 or Myh10 alone, or both simultaneously, using the PDGFB-CreERT2 line (24). Single deletion of Myh10 in endothelial cells (MIIB EC-KO) did not significantly affect retinal vascular morphogenesis, although we noticed a small decrease in the number of filopodia in endothelial tip cells (Sup.Fig.3A-C). Efficient depletion of MIIB was observed by immunofluorescence (Sup.Fig.3B). Single deletion of Myh9 (MIIA EC-KO) showed a very distinct phenotype. MIIA EC-KO retinas showed a mild decrease in radial expansion, an increase in vessel density, decreased proliferation and an increase in the number of tip cells (Fig.1D-F). In accordance with a mild radial expansion phenotype, vascularization of the superficial in MIIA EC-KO animals reached the retina margin by P12 (Sup.Fig.4). Yet, the most striking phenotype is a dramatic change in tip cell morphology (Fig.1E, and Sup.Fig.4A). MIIA-deficient tip cells showed an abnormal branched morphology with extremely long and enlarged membrane protrusions and severe decrease in filopodia number (Fig.1E, and Sup.Fig.5A). Remarkably, the absence of filopodia and enlarged membrane protrusions was tip cell-specific, as endothelial cells in non-sprouting areas showed numerous filopodia, morphologically similar to WT control littermates, even if MIIA-deficient endothelial cells in the plexus show a trend to have a decreased number of filopodia (Sup.Fig.5B,C). Immunofluorescence analysis confirmed the efficient depletion of MIIA (Fig.1G), and that the MIIA EC-KO phenotype manifested even in the presence of high-levels of MIIB in tip cells (Sup.Fig.5D). Despite the notable changes in tip cell morphology, filamentous actin did not appear to be severely compromised, with noticeable actin cables present in the cell cortex (Fig.1G).

**Figure 1:**
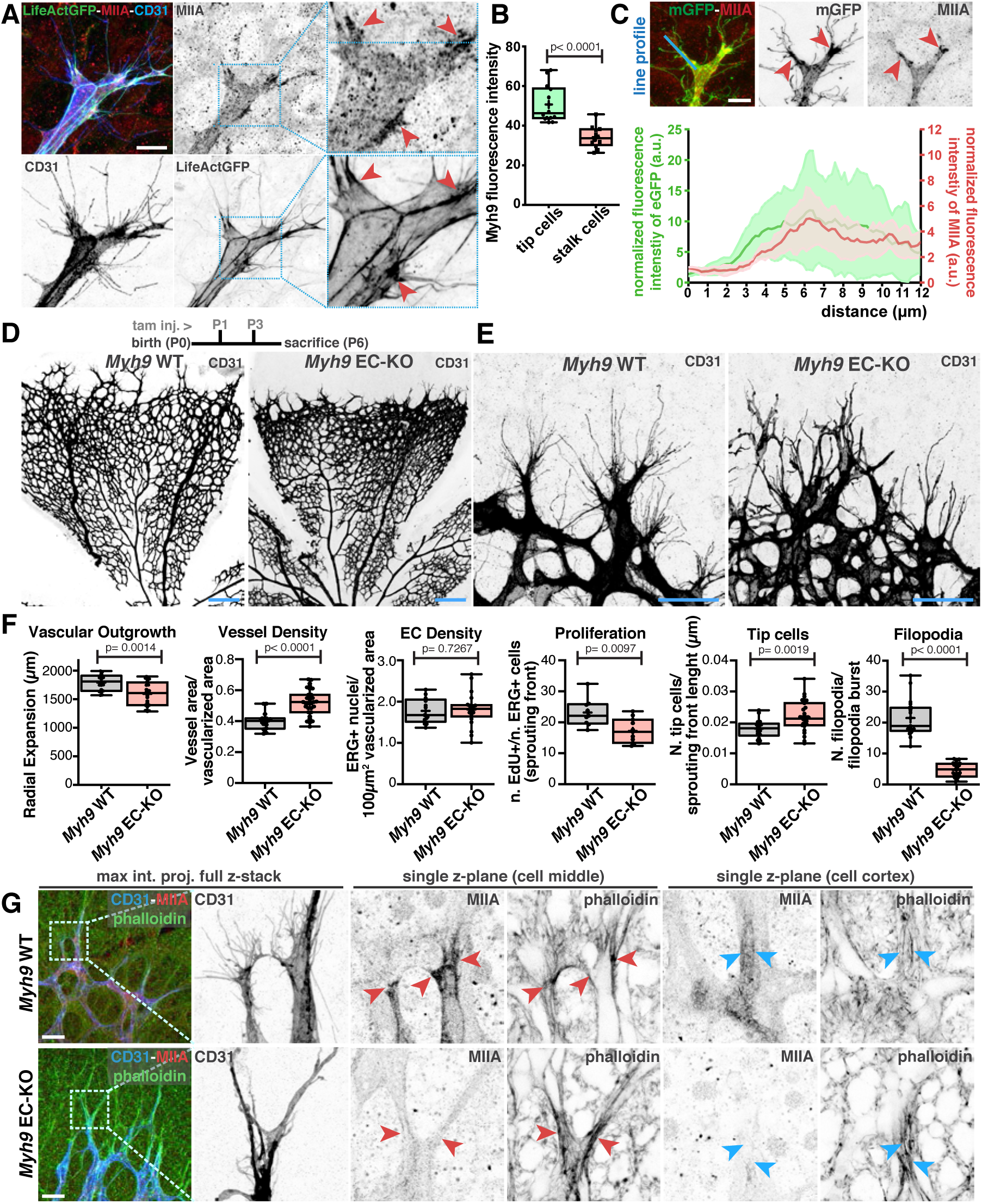
(A) Representative images of tip cells from LifeAct-GFP mouse retinas labeled for MIIA (red), actin (green) and CD31 (blue). Red arrowheads point to sites enriched in MIIA at the base of filopodia. Scale bar, 20 µm. (B) Box plot of Myh9 fluorescence intensity between tip and stalk cells from PDGFB-iCRE:mTmG mouse retinas (n = 14 tip cells, n = 12 stalk cells). *p*-value from unpaired student t-test. (C) Representative image (upper panel) and graph (bottom panel) of the normalized line-scan fluorescence intensity profile (blue line) for MIIA and mGFP signals from endothelial tip cell’s leading edge (n = 6 endothelial tip cells). Red arrowheads point to sites enriched in MIIA at the base of filopodia. Scale bar, 20 µm. (D) Upper panel: timeline of tamoxifen injection (tam. inj.) and age of sacrifice of mouse pups. Representative images of mouse retinas from *Myh9* WT and *Myh9* EC-KO labeled for CD31. Scale bar, 250 µm. (E) Representative images of tip cells and filopodia in the angiogenic sprouting front of mouse retinas from *Myh9* WT and *Myh9* EC-KO labeled for CD31. Scale bar, 50 µm. (F) Box plot of vascular outgrowth, vessel density, EC density, EC proliferation, number of tip cells and number of filopodia per filopodia burst in *Myh9* WT (n = 19 retinas) and *Myh9* EC-KO (n = 24 retinas) mouse retinas. *p*-value from unpaired student t-test. (G) Left panel: high-magnification images of tip cells labeled for CD31 (blue), F-actin (green) and MIIA (red) from *Myh9* WT and *Myh9* EC-KO. Right panel: higher-magnification images from delineated regions in left panel for single z-plane of tip cells in two different planes. Red arrowheads points towards leading edges of tip cells, blue arrowheads points towards cortical actin cables in tip cells. Scale bar, 50 µm.

Abrogation of both Myh9 and Myh10 led to an extremely severe phenotype, where endothelial cells at the sprouting front lacked filopodia and showed a very elongated and slim membrane protrusions, likely caused by a complete disruption of actomyosin contractility (Sup.Fig.6A-C). Altogether, these results revealed that MIIA plays a fundamental role in the formation of tip cell’s filopodia, which cannot be rescued by MIIB. In addition, MIIA and MIIB, together, are essential to maintain endothelial cell shape. This is consistent with previous observation that both MII isoforms play overlapping but also unique roles (23, 25–27).

The specific inhibition of filopodia formation and the presence of long membrane protrusions in MIIA-deficient tip cells, suggested that MIIA plays an important role in the regulation on the type of membrane protrusions that tip cells produce (Fig.1E, G, and Sup.Fig.5A). Thus, we decided to investigate the dynamics of membrane protrusions in endothelial tip cells *in vivo*. To do so, we established a new protocol for live-imaging of the mouse retina (detailed description in material & methods). Live-imaging showed that WT endothelial tip cells have a high rate of filopodia formation (∼0.6 filopodia/sec) (Fig.2A,B; Sup.Fig.7A,B and Sup.Videos 3-6).

**Figure 2:**
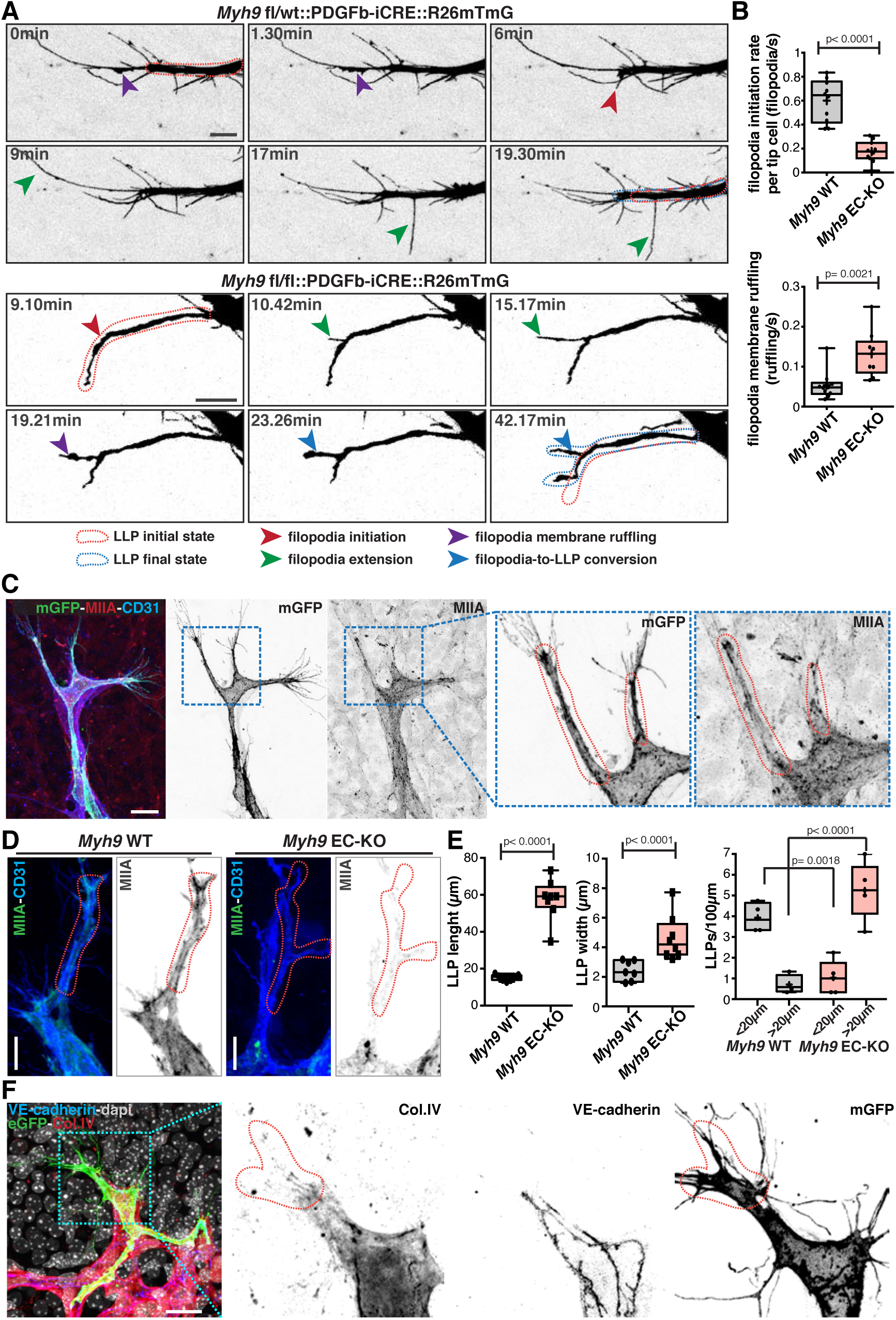
(A) Time-course still images of LLPs and filopodia dynamics in *Myh9 fl/wt::PDGFB-iCRE::R26mTmG* and *Myh9fl/fl::PDGFB-iCRE::R26mTmG* mouse retinas. Red dashed line contour represents LLPs initial state. Blue dashed line contour represents LLPs final state. Red arrow indicates filopodia initiation; green arrow indicates filopodia extension; purple arrow indicates filopodia membrane ruffling; and blue arrow indicates filopodia-to-LLPs conversion. Scale bar, 10 µm. (B) Box plot of filopodia initiation rate per tip cell (filopodia/s) (*Myh9* WT n = 10 tip cells; *Myh9* EC-KO n = 11 tip cells) and filopodia membrane ruffling (ruffling/s) in *Myh9* WT (*Myh9* WT n = 10 filopodia) and *Myh9* EC-KO (*Myh9* EC-KO n = 9 filopodia) mouse retinas. *p*-value from unpaired student t-test. (C) Representative images of a tip cell from *PDGFb-iCre::R26mTmG* mouse retina labeled for MIIA (red), cell membrane (green), CD31 (blue), highlighting LLPs in tip cells (red dashed line), with numerous filopodia in their extremity. Scale bar, 20 µm. (D) Representative images of tip cells and LLPs from *Myh9* WT and *Myh9* EC-KO mouse retinas labeled for MIIA (green) and CD31 (blue). Scale bar, 10 µm. Red dashed line contours LLPs. (E) Box plot of number of LLPs (<20µm or >20µm) in *Myh9* WT (n = 4 retinas) and *Myh9* EC-KO mouse retinas (n = 5 retinas), and LLP width and length in *Myh9* WT (n = 7 retinas) and *Myh9* EC-KO mouse retinas (n = 8 retinas). *p*-value from unpaired student t-test. (F) Representative images of LLPs in tip cells from PDGFb-iCRE::R26mTmG (green) a P6 mouse retina labeled for VE-cadherin (blue), Col.IV (red), and nuclei (dapi, grey). Scale bar, 20 µm. Red dashed line contours LLPs.

Surprisingly, live-imaging of MIIA-deficient retinas showed that tip cells were also able to protrude filopodia, albeit with significant altered dynamics (lower rate of initiation; lower extension speed, and duration of extension) (Fig.2A,B, Sup.Fig.7A,B and Sup.Videos 7-11). Intriguingly, in both MIIA WT and MIIA EC-KO tip cells, we observed an unexpected membrane ruffling activity at the filopodia membrane (Fig.2A, Sup.Fig.7A and Videos 7-11). WT tip cells exhibited membrane ruffling both at filopodium’s base and along filopodium’s lateral membrane (Fig.2A and Sup.Videos 3-6), yet, ruffling activity was not a general feature of all filopodia. In our imaging conditions, only a small proportion of filopodia showed ruffling activity in their lifespan, and most of them extended and retracted without detectable ruffling activity. Moreover, when present, filopodium’s ruffling was normally short-lived (1 to 6 minutes). Ruffling activity at filopodium’s base was associated with extension and enlargement of the cytoplasmic membrane (Fig.2A and Sup.Fig.7A,B). Membrane ruffling is highly associated to actin polymerisation. To confirm this link, we live-imaged WT LifeAct-GFP retinas that showed high and dynamic actin polymerization at the base of filopodia (Sup.Fig.7C, and Sup.Videos 12 and 13), suggesting that membrane ruffling is indeed driven by actin polymerisation. In contrast to MIIA WT tip cells, in MIIA EC-KO tip cells, membrane ruffling was more persistent and generally led to the conversion of filopodia into long membrane protrusions, similar to the ones observed in fixed samples (Fig.2A, Sup.Fig.7A and Videos 7-11). Thus, our live-imaging observations indicate that long membrane protrusions in MIIA-deficient tip cells are originated from altered filopodia dynamics, through excessive actin-dependent membrane ruffling. Moreover, they suggest that one key function of MIIA in tip cells is to negatively regulate the formation of such long protrusions.

Given that MIIA WT tip cells also showed substantial filopodia membrane ruffling, we reasoned that the formation of long membrane protrusions might be a particular feature of WT tip cells that becomes overrepresented in MIIA EC-KO. Indeed, careful analysis of WT tip cells showed similar morphological membrane protrusions, comparable to the prominent ones observed in MIIA EC-KO tip cells, yet their frequency and size were much reduced (Fig.2C-E and Sup.Fig.8A). In WT tip cells, these membrane protrusions were characterized by a mean length of ∼20µm (ranging from ∼5 to ∼33 µm) and a mean width of ∼2µm (ranging from ∼0.7 to ∼5µm) (Fig.2E), enriched in actin and MIIA, and from which numerous filopodia emanated (Fig.2C,D and Sup.Fig.8B). Given their morphology and link to membrane-ruffling activity, we named these naturally occurring protrusions as long lamellipodia projections (LLPs). Interestingly, the presence of LLPs correlated with regions where endothelial tip cells contacted with non-vascular extracellular matrices (ECM), highlighted by decreased staining of the vascular basement membrane marker collagen IV (Fig.2F and Sup.Fig.8C). Moreover, LLPs were associated with increased levels of active integrin beta 1 (ITGB1), a marker for matrix-bound focal adhesions (Sup.Fig.8C). LLPs in MIIA-deficient endothelial cells also correlated with sites of contact with non-vascular extracellular matrix and with activated ITGB1, however its expression pattern appeared to be more diffuse along LLPs and not enriched at the base of filopodia, as in WT cells (Sup.Fig.8C). Altogether, these data suggest that LLPs are endothelial tip cell-specific protrusions derived from filopodia that might play roles in invasion and migration into non-vascular ECM.

Next, we investigated the mechanism driving LLP formation in endothelial cells. Actin-based membrane ruffling is known to rely on Arp2/3-dependent dendritic actin networks (6). To confirm if Arp2/3 is involved in LLP formation, we specifically deleted Arpc4, a structurally essential component of the Arp2/3 complex (5, 28), in endothelial cells *in vivo*. Deletion of Arpc4 in endothelial cells, Arpc4 EC-KO, led to a very prominent vascular phenotype. Endothelial cells without a functional Arp2/3 complex were unable to invade into avascular areas, leading to a dramatic reduction in radial expansion at P6 (Fig.3A-C). Higher magnification of tip cells demonstrated that Arpc4-deficient cells had significantly impaired LLPs formation, which correlated with an increase in the number and length of filopodia (Fig.3B,D,E). Strikingly, Arpc4 EC-KO phenotype inversely mirrors MIIA EC-KO phenotype in terms of tip cell’s filopodia and LLPs number and morphology (compare Fig.1E,F and Fig.3B,C). Yet, Arp2/3 deficiency did not alter MIIA distribution in endothelial tip cells (Fig.3D).

**Figure 3:**
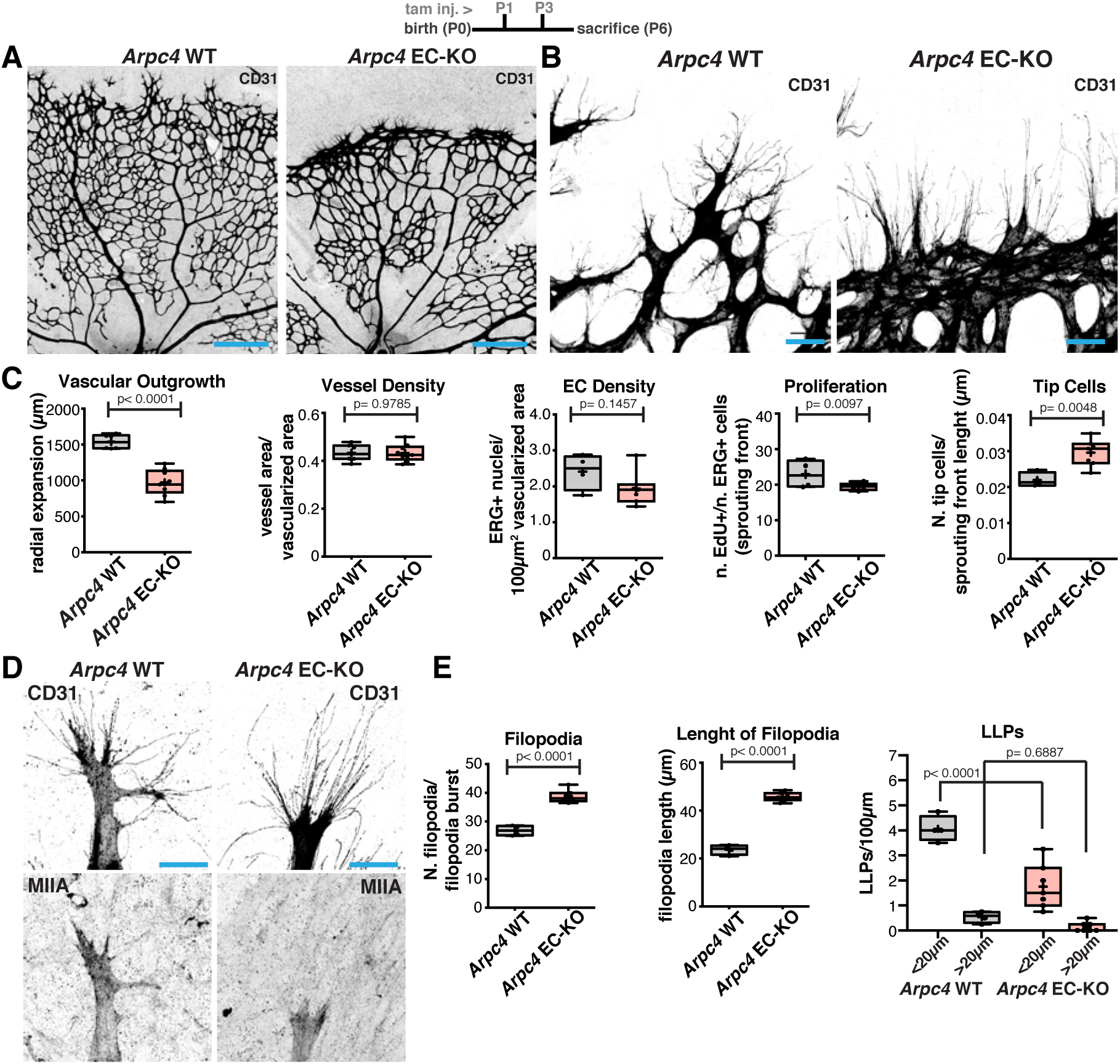
(A) Representative images of mouse retinas from *Arpc4* WT and *Arpc4* EC-KO labeled for CD31. Scale bar, 250 µm. (B) Representative images of tip cells and filopodia in the angiogenic sprouting front of mouse retinas from *Arpc4* WT and *Arpc4* EC-KO labeled for CD31. Scale bar, 50 µm. (C) Box plot of vascular outgrowth, vessel density, EC density, EC proliferation and number of tip cells per sprouting front length (µm) in *Arpc4* WT (n = 7 retinas) and *Arpc4* EC-KO (n = 9 retinas) mouse retinas. *p*-values from unpaired student t-test. (D) Representative images of tip cells from *Arpc4* WT and *Arpc4* EC-KO mouse retinas labeled for CD31 and MIIA. Scale bar, 20 µm. (E) Box plot of number of filopodia per filopodia burst, length of filopodia (µm) and number of LLPs per 100µm (<20µm or >20µm) in *Arcp4* WT (n = 4 retinas) and *Arpc4* EC-KO (n = 7 retinas) mouse retinas. *p*-values from unpaired student t-test.

**Figure 4:**
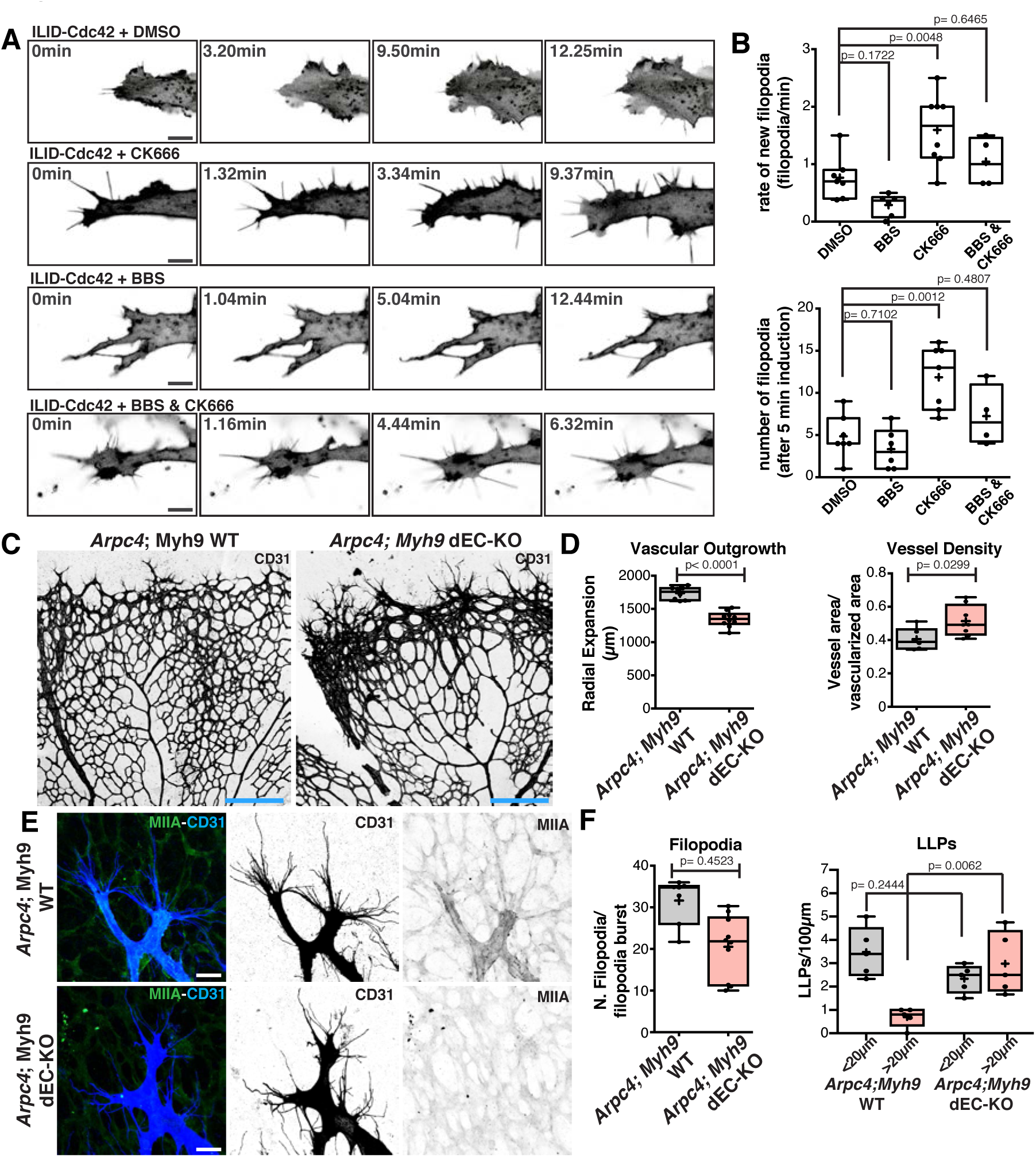
(A) Representative images of HUVECs expressing an optogenetic activator of Cdc42 (ILID) in DMSO, BBS, CK666, and BBS + CK666 conditions. Scale bar, 10 µm. (B) Box plot of rate of new filopodia and total number of filopodia in different conditions (DMSO n = 7 cells; BBS n = 6 cells; CK666 n = 8 cells; and BBS + CK666 n = 4 cells). *p*-values from unpaired ANOVA test. (C) Representative images of mouse retinas from *Arpc4;Myh9* WT and *Arpc4*;*Myh9* dEC-KO mouse retinas labeled for CD31. Scale bar, 250 µm. (D) Box plot of vascular outgrowth and vessel density in *Arpc4;Myh9* WT (n = 8 retinas) and *Arpc4*;*Myh9* dEC-KO (n = 10 retinas) mouse retinas. *p*-values from unpaired student t-test. (E) Representative images of tip cells and filopodia/LLPs from *Arpc4;Myh9* WT and *Arpc4*;*Myh9* dEC-KO mouse retinas labeled for MIIA (green) and CD31 (blue). Scale bar, 20 µm. (F) Box plot of number of filopodia per filopodia burst and number of LLPs (<20µm or >20µm) in *Arpc4;Myh9* WT (n = 5 retinas) and *Arpc4*;*Myh9* dEC-KO mouse retinas (n = 5 retinas). *p*-value from unpaired Student t-test.

Analysis of P12 retinas showed that radial expansion was completely abrogated, as radial outgrowth was mostly at the same distance from the optic nerve as in P6 retinas (Sup.Fig.9A). Altogether, we conclude that LLPs are formed by Arp2/3-dependent actin polymerisation and that endothelial tip cell invasiveness is entirely determined by Arp2/3 activity. Moreover, it proves that presence of filopodia is insufficient for tip cell invasion.

Next, we investigated how the balance between filopodia and LLPs in endothelial tip cells is established. Remarkably, MIIA KO phenotype was characterized by a loss of filopodia and a gain in LLPs, whilst Arpc4 EC-KO displayed excessive filopodia and reduced number of LLPs. Moreover, we showed that LLPs originated from filopodia. Thus, we hypothesized that MIIA could balance the ability of tip cells to form filopodia over LLPs, by regulating Arp2/3 complex activity. A prediction from this model is that inhibition of Arp2/3 complex in MIIA-deficient cells should rescue the ability of endothelial cells to produce filopodia. To test this hypothesis, we turned to *in vitro* human umbilical vein endothelial cell (HUVEC) cultures. Endothelial cells growth on glass or plastic did not show the ability to form long filopodia. However, when seeded on top of fibroblast monolayers, HUVECs acquired this capacity (29). However, in both *in vitro* conditions, HUVECs did not display protrusions similar to LLPs, but rather standard lamellipodia. DMSO-treated or CK666-treated (a specific inhibitor of Arp2/3) cells have similar number of filopodia in normal conditions. In contrast, blebbistatin (BBS) treatment, an inhibitor of MII activity, abrogated filopodia formation. Remarkably, filopodia formation in BBS-treated cells was partially rescued by co-treatment with CK666 (Sup.Fig.10A,B). These results were further confirmed by siRNA-mediated knock-down of MIIA and Arp2/3 complex (Sup.Fig.10C,D). To further confirm these observations, we used an optogenetic tool that allows timed and local activation of Cdc42 (30), the main regulator of endothelial cell protrusions *in vivo* and *in vitro* (31). We observed that in DMSO-treated conditions, Cdc42 activation led to high lamellipodia activity and few filopodia, which was reverted by an inhibitor of Arp2/3 (CK666) (Fig.5A,B and Sup.Video 14 and 15). In contrast, BBS-treatment abrogated filopodia formation (Fig.5A,B and Sup.Video 16). The capacity of forming filopodia in BBS-treated cells was restored by co-treatment with CK666 (Fig.5A,B and Sup.Video 17), supporting the hypothesis that MII-activity promotes filopodia formation by limiting the activation of Arp2/3. To verify if MIIA EC-KO leads to a lack of filopodia and excessive LLPs due to excessive Arp2/3 activity *in vivo*, we generated a double loss-of-function of both Arp2/3 and MIIA, by crossing MIIA EC-KO and Arpc4 EC-KO mice. Overall, Arpc4::MIIA EC double KO mice showed a phenotype very similar to Arpc4 EC-KO, with a strong reduction in radial expansion and compaction at the vascular front (Fig.5C,D). Yet, abrogation of Arp2/3 rescued filopodia formation in MIIA-deficient endothelial cells *in vivo* (compare Fig.5E,F, Fig.1E,F, and Fig.2E). Altogether, these results demonstrate *i)* that Arp2/3 activity is necessary for endothelial tip cell invasiveness and formation of pro-invasive LLPs; *ii)* that LLPs derive from filopodia; *iii)* that filopodia are not required for tip cell migration; *iv)* and that MIIA enables filopodia formation by restricting Arp2/3 activation in endothelial tip cells *in vivo*; .

**Figure 5:**
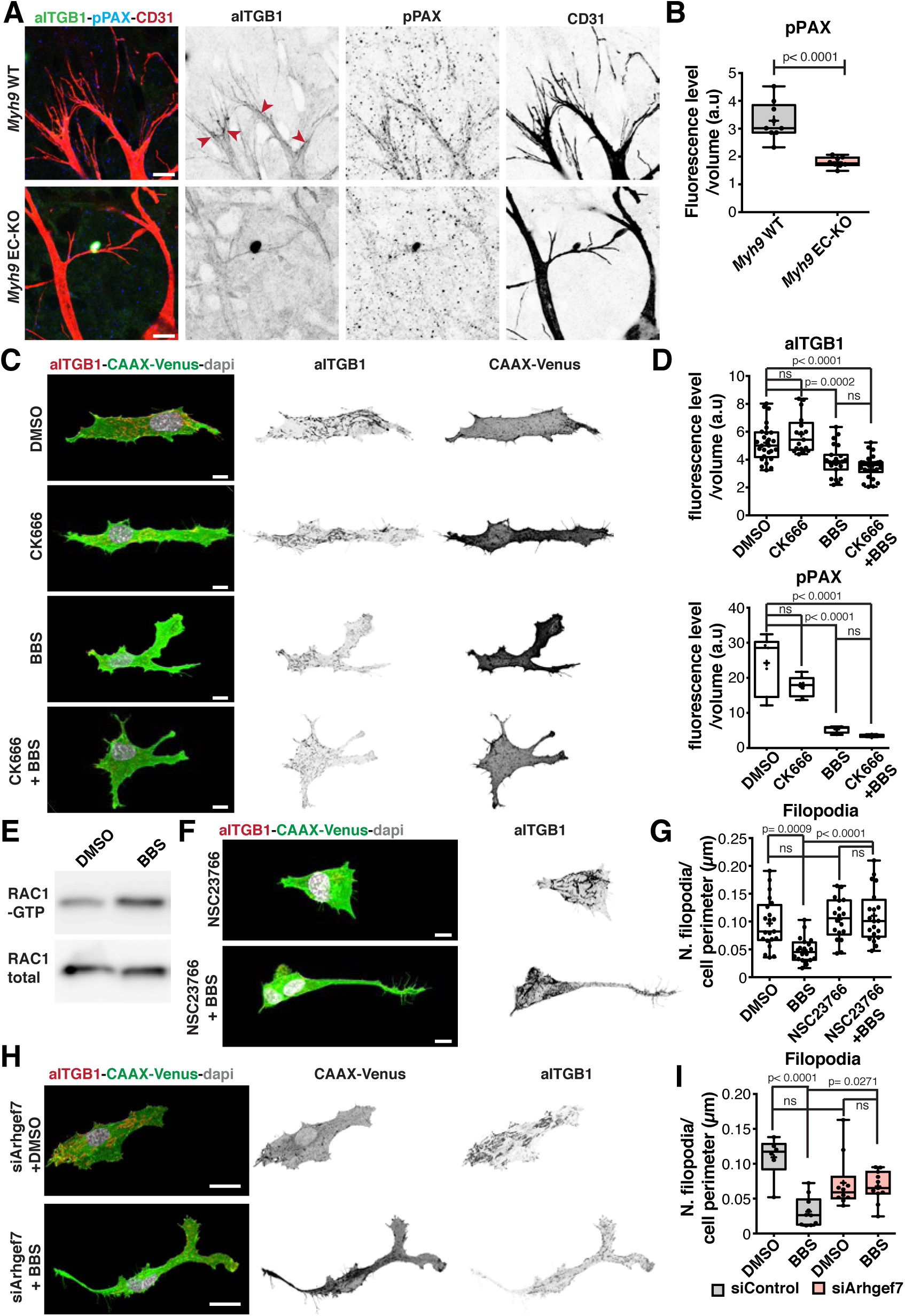
(A) Representative images of tip cells from Myh9 WT and *Myh9* EC-KO mouse retinas labeled for pPax Y118 (blue), activated ITGB1 (aITGB1, green) and CD31 (red). Scale bar, 10 µm. Red arrowheads point to sites enriched in aITGB1 and pPAX at the base of filopodia in *Myh9* WT tip cells, which is absent or more homogenously distributed along LLPs of *Myh9* EC-KO tip cells. Scale bar, 20 µm. (B) Box plot of pPax fluorescence in tip cells from *Myh9* WT (n = 9 retinas), *Myh9* EC-KO (n = 8 retinas) mouse retinas. *p*-value from unpaired student t-test. (C) Representative images of HUVECs expressing CAAX-Venus at the cell membrane in DMSO, BBS, CK666 and BBS + CK666 conditions. HUVECs labeled for activated ITGB1 (red), cell membrane (CAAX-Venus, in green) and nuclei (DAPI, in grey). Scale bar, 10 µm. (D) Box plot of pPax fluorescence intensity and activated ITGB1 fluorescence intensity in HUVECs expressing CAAX-Venus at the cell membrane in DMSO (n = 20 cells), BBS (n = 15 cells), CK666 (n = 24 cells) and BBS + CK666 (n = 13 cells) conditions. *p*-values from one-way ANOVA. (E) Western blot analysis of levels of active GTP-bound Rac1 and total Rac1 in DMSO or BBS-treated HUVECs. (F) Representative images of HUVECs expressing CAAX-Venus at the cell membrane in NSC23766 and NSC23766 + BBS conditions. HUVECs labeled for activated ITGB1 (red), cell membrane (CAAX-Venus, in green) and nuclei (DAPI, in grey). Scale bar, 10 µm. (G) Box plot of filopodia number in DMSO (n = 21 cells), BBS (n = 20 cells), NSC23766 (n = 18 cells), and NSC23766 + BBS (n = 20 cells). *p*-values from one-way ANOVA. (H) Representative images of HUVECs expressing CAAX-Venus at the cell membrane in siArhgef7 HUVECs treated with DMSO or BBS. HUVECs labeled for activated ITGB1 (red), cell membrane (CAAX-Venus, in green) and nuclei (DAPI, in grey). Scale bar, 10 µm. (I) Box plot of filopodia number in siControl and siArhgef7 HUVECs treated with DMSO (n = 6 cells in siControl, and 10 in siArhgef7) or BBS (n = 10 cells in siControl, and 11 in siArhgef7). *p*-values from one-way ANOVA.

We next investigated the mechanism by which MIIA limits Arp2/3 activity. Arp2/3 is activated by WASP/WAVE complexes downstream of the Rho GTPases Rac1 and Cdc42 (32). A common trigger for Rac1/Cdc42 activation and migration is signalling from immature FAs at the leading edge of cells (32, 33). Our observations showed that Arp2/3-dependent LLPs correlated with local invasion into extravascular matrices (Fig.2F, Sup.Fig.7), suggesting that integrin signalling from focal adhesions could activate Arp2/3 when engaging with extravascular matrices. In accordance, ITGB1 EC-KO (34) shows striking similarities with Arpc4 EC-KO, including excessive and longer filopodia formation, blunted sprouting front (reminiscent of a lack of LLPs), and severely reduced invasion to the deeper layers. Moreover, it is well established that MII-dependent activity promotes maturation of integrin adhesions (35). Thus, we hypothesized that FAs could be platform promoting the crosstalk between MIIA and Arp2/3 at the leading edge of tip cells. To evaluate the state of FAs, we analysed integrins (ITGA5 and activated ITGB1) and phosphorylated paxillin (pPAX, a marker for more mature focal adhesions) at the base of filopodia and in LLPs in endothelial tip cells (Fig.6A and Sup.Fig.11A,B). MIIA-deficient tip cells showed a strong and significant reduction of pPAX, and signals for ITGA5 and activated ITGB1 were not enriched at the base of filopodia and diffused all over LLPs (Fig.6A,B and Sup.Fig.11A,B), suggesting that inhibition of MIIA leads to significant differences in integrin location and activation state. Similar results were observed in endothelial cells *in vitro*. BBS-mediated inhibition of MII in HUVEC-fibroblasts co-cultures led to a strong decrease in the number of mature fibrillar FAs and a strong decrease in pPAX, both at immunofluorescence and western blot level, whilst CK666-treatment did not modify levels of pPAX (Fig.6C,D and Sup.Fig.12A-C), which correlates with a decrease in number of filopodia (Sup.Fig.10). Our results are suggestive of a mechanism where immature/nascent adhesions would activate Arp2/3, and MIIA inhibits this signalling axis by promoting maturation of FAs, as previously reported (35, 36). We next assessed if BBS treatment would promote activation of Rac1, a major positive regulator of Arp2/3 in endothelial cells downstream of integrin signalling. We observed that BBS treatment increased levels of Rac1-GTP, the active form of Rac1, when compared to DMSO treatment (Fig.6E). Accordingly, inhibition of Rac1 activation with NSC23766 led to a rescue of filopodia formation in BBS-treated cells (Fig.6G and Sup.Fig.13). Tiam1, DOCK180, and Arhgef7/βPIX were previously implicated in Rac1 activation downstream of integrins (37). In addition, previous work showed that MIIA inhibits βPIX recruitment to and activation by focal adhesions (36). Thus, we tested if knockdown of βPIX could rescue filopodia formation in BBS-treated cells. Indeed, silencing of βPIX (Sup.Fig.14A) led to a partial rescue in filopodia number in BBS-treated endothelial cells (Fig.6H-I and Sup.Fig.14B). Taken together, these results suggest that MIIA balances LLPs and filopodia formation by promoting FA maturation and thereby limiting Arp2/3 activation, through inhibition of Rac1 activation downstream of integrin-βPIX in nascent adhesions.

## Discussion

Deciphering the mechanisms used by endothelial cells to migrate and invade is essential to understand sprouting angiogenesis and to develop novel anti-angiogenic therapies. We found that invasiveness of endothelial tip cells during angiogenesis *in vivo*, depends on the formation of a specific Arp2/3-dependent protrusion, the LLP. We found that LLPs are derived from filopodia at sites in contact with non-vascular extracellular matrix, and enable efficient invasion of endothelial cells *in vivo*. Our findings also support the idea that filopodia are not essential for endothelial cell invasion and migration, as previously reported (21).

Previous studies have highlighted that Arp2/3 and formins compete with each other to regulate the organization of filamentous actin and to dictate the type of cellular protrusions (38, 39). Remarkably, we show for the first time that MIIA balances the relative proportion of two types of cellular protrusions, favoring filopodia over LLP, a lamellipodia-like structure. This is rather surprising as increased contractility is generally inversely correlated with protrusive activity, as shown in several different models (9, 10). Moreover, we discovered that MIIA balances the protrusive activity of tip cells by limiting Arp2/3 activation through actomyosin contractility-induced maturation of focal adhesions. Notably, this effect was restricted to MIIA and not MIIB, in agreement with reports demonstrating that MII isoforms have unique roles (23, 25–27).

Based on this study and on previous reports, we propose an integrative view on the mechanisms of endothelial tip cell’s invasive behaviour: VEGFA controls formins, by directly regulating Cdc42-activity (31, 40) and MIIA, via SRF transcriptional activity (16). This fosters actomyosin contractility and protrusive activity in endothelial tip cells. Cdc42/formin-dependent filopodia activity will engage with extravascular matrices, activating integrin signalling at the base and within filopodia, which promotes Arp2/3 activity downstream of a beta-PIX/Rac1 pathway. MIIA positively regulates the maturation state of focal adhesions, which negatively regulates the βPIX/Rac1 pathway, and thus prevents excessive Arp2/3 activation. Loss of MIIA activity leads to overactivation of Arp2/3 and unrestrained conversion of filopodia into LLPs. Loss of Arp2/3 activity abrogates LLPs formation and thus inhibits cell migration and invasion (Sup.Fig.15). Excessive filopodia formation in Arp2/3-defficient endothelium can be explained by the previously documented competition between actin nucleators (41). Within this model, an outstanding question relates to the mechanism of local control of MIIA activity that licenses LLP formation. We can speculate that filopodia-dependent signalling (engagement with local guidance cues, extravascular matrix components or cells) would trigger inhibition of MIIA at the base of filopodia leading to Arp2/3 activation and LLPs formation. Exploration of this molecular crosstalk might provide new therapeutic opportunities to sprouting angiogenesis based on the ability to block invasion of endothelial tip cells.

## MATERIALS AND METHODS

### Mice and treatments

In this study, we used the following mouse strains: *Myh9* floxed (42); *Myh10* floxed (43) *Arpc4* floxed (28); LifeAct.GFP (44); R26mTmG (45); Myh9.GFP (46); *Pdgfb*-iCreERT2 (24). Intercrosses between the different mouse strains generated new double and triple transgenic mouse strains. As controls, Cre-negative littermates were used in all experiments. Both males and females were used, without distinction. Tamoxifen (Sigma, Germany) was injected intraperitoneally (IP) (20 µl/g of 1 mg/mL solution) at postnatal day 1 (P1) and P3 before eyes were collected either at P6 and P12 or injected at P4 and P5 and collected at P12, as described previously (47).

Mice were maintained at the Instituto de Medicina Molecular (iMM) under standard husbandry conditions and under national regulations, with the exception of Myh9.GFP which was kept in the Institut Curie Specific Pathogen Free (SPF) animal facility for breeding. Animal procedures were performed under the DGAV project license 0421/000/000/2016.

### Immunofluorescence on mouse retinas

Eyes were collected at P6 and at P12 and fixed with 2% PFA in PBS for 5 hr at 4°C, thereafter retinas were dissected in PBS. Blocking/permeabilization was performed using CBB (16), consisting of 1% FBS (Thermo Fisher Scientific), 3% BSA (Nzytech), 0.5% triton X100 (Sigma), 0.01% Na deoxycholate (Sigma), 0,02% Na Azide (Sigma) in PBS pH = 7.4 for 2 hr in a rocking platform. Primary antibodies used: anti-CD31 (R&D, AF3628, 1:200), anti-Erg (Abcam, ab92513, 1:200), anti-Col.IV (BioRad, 2150-1470, 1:400), anti-MIIA (SIGMA, M8064, <u>1:600</u>), anti-MIIB (SIGMA, M7939, <u>1:400</u>), anti-aITGB1 (BD Pharmingen, 553715, <u>1:100</u>), anti-pPaxY118 (Cell Signalling, 2541S, 1:100), anti-bPix (Merck, 07-1450-I, <u>1:100</u>), anti-pMLC2 (Abcam, ab2480, 1:200), anti-ITGA5 (Abcam, ab150361, 1:100) were incubated in 1:1 CBB:PBS at 4°C overnight in a rocking platform and afterwards washed 3 × 60 min in PBS-T. Then, retinas were incubated in 1:1 CBB:PBS solution containing the secondary fluorophore-conjugated antibodies at 4°C overnight in the dark. The secondary antibodies used were: Donkey anti-goat Alexa 488 (Thermo Fisher Scientific, A11055, 1:400), Alexa 555 (Thermo Fisher Scientific, A21432, 1:400), Alexa 647 (Thermo Fisher Scientific, A21447, 1:400), Donkey anti-rabbit Alexa 488 (Thermo Fisher Scientific, A21206, 1:400), Alexa 568 (Thermo Fisher Scientific, A10042, 1:400), Alexa 647 (Thermo Fisher Scientific, A31573, 1:400), Donkey anti-rat Alexa 555 (Thermo Fisher Scientific, A21208, 1:400), Goat anti-rat Alexa 555 (Thermo Fisher Scientific, A21434, 1:400) and Phalloidin Alexa 488 (Thermo Fisher Scientific, A12379, 1:400) or Phalloidin Alexa 555 (Thermo Fisher Scientific, A12380, 1:400). Retinas were mounted on slides using Vectashield mounting medium (Vector Labs, H-1000, Burlingame, California, USA). For vascular parameters quantification, representative images were acquired on a Zeiss Cell Observer Spinning Disk microscope, equipped with the Zen software with a Plan-Apochromat 40x/1.4 Oil DIC M27 objective or in a confocal Laser Point-Scanning Microscope 880 (Zeiss) equipped with the Zen black software with a Plan Apochromat 20x NA 0.80 dry objective, a C-Apochromat 40x NA 1.20 Water objective or a 63x NA 1.40 oil DIC M27 objective.

### Retina live-imaging

Eyes were collected from P6 mouse pups and dissected fresh in warm Leibovitz’s L-15 Medium (Thermo Fisher, 21083027) with 2% FBS (Thermo Fisher, 10500064), flat-mounted embedded in 0.5% low-melting agarose (VWR) into a 60mm cell culture petri dish. Retinas were then live-imaged using a TCS SP8 MP microscope (Leica) at 900nm wavelength (Spectra Physics InSight DS+ Dual) using a water immersion objective (Leica, 25x, 0.95 NA) warmed at 37°C.

### Culture of HUVECs and NIH3T3 Fibroblasts

NIH3T3 fibroblasts (ATCC) were routinely grown in DMEM medium supplemented with 10% fetal bovine serum (Thermo Fisher, 10500064) and 0.01% Penicillin/Streptomycin (Thermo Fisher, 15140122). Cells were maintained in 37 °C humidified atmosphere with 5% CO2. Human umbilical vein endothelial cells – HUVECs (C2519A, Lonza) – were routinely cultured following the manufacturer’s guidelines, in filter-cap T75 flasks Nunclon Δ surface treatment (VWR international, LLC) with complete medium EGM-2 Bulletkit (CC-3162, Lonza) supplemented with 0.01% Penicillin/Streptomycin (#15140122, Gibco) at 37°C and 5% CO_2_ to ensure a stable environment for optimal cell growth. All experiments were performed with HUVECs between passages 3 and 6.

When passaging HUVECs for experiments, cells were washed twice in sterile PBS (137mM NaCl, 2.7mM KCl, 4.3mM Na_2_HPO_4_, 1.47mM KH_2_PO_4_, pH7.4) and then incubated for 2-3min in Trypsin/EDTA (#15400054, Gibco) at 37°C, 5% CO_2_. When 95% of the cells detached, complete medium was added to each flask to inhibit the activity of the Trypsin/EDTA and the cell suspension was transferred onto a falcon tube. To maximize the amount of cells collected, all the flasks were washed again with complete medium, which was added to the cell suspension gathered previously. Cells were then centrifuged at 700rpm for 5min at RT and the pellet re-suspended in fresh complete medium. The cell concentration present in the suspension was determined using a Neubauer Chamber Cell Counting (Hirschmann EM Techcolor). HUVECs were then seeded on the desired culture vessels at 210.000–300.000 cells/mL and placed in the incubator.

### siRNA transfection

In order to silence the expression of genes of interest, a set of ON-TARGET human siRNAs against Arp3 (J-012077-08, Dharmacon, GE Healthcare), Myh9 (J-007668-07, Dharmacon GE Healthcare), beta-Pix/ArghGEF7 (J-009616-07, Dharmacon GE Healthcare) or untargetting control were used. Briefly, HUVECs were seeded the day before the transfection to reach 60-70% confluence and were then transfected with 25 nM of siRNA using the DharmaFECT 1 reagent (Dharmacon, GE Healthcare) following the Dharmacon siRNA transfection protocol. 24h after transfection the culture medium was replaced by fresh complete medium and cells were kept under culture conditions up until 72h post-transfection and then processed for live-imaging, immunofluorescence, protein or RNA extraction.

### Viral production and transduction

Replication-incompetent lentiviruses were produced by transient transfection of HEK293T with lentiviral expression vector co-transfected with the viral packaging vector Δ8.9 and the viral envelope vector VSVG. Medium was replaced with fresh culture medium 6-8h post transfection. 48h after medium replacement, lentiviral particles were concentrated from supernatant by ultracentrifugation at 112.500g for 1h30 and re-suspended in 0.1% BSA PBS. Seeded HUVECs were transduced 24h post-transfection with varying concentrations of lentiviral plasmids containing CAAX-Venus (pLL7.0: Venus-iLID-CAAX (from KRas4B); Addgene plasmid # 60411) and/or ILID-Cdc42 (pLL7.0: hITSN1(1159–1509)-tgRFPt-SSPB WT; Addgene plasmid # 60413) were gifts from Brian Kuhlman (30). 24h after viral transduction the culture medium was replaced by fresh complete medium and cells were kept under culture conditions up until 48h post-transduction and then processed for immunofluorescence or imaging.

### HUVECs-fibroblasts co-cultures

For HUVECs and fibroblasts co-cultures, the fibroblasts were first seeded at concentration of 150,000 cells/well on 24-well plates with glass coverslips, previously coated with 0.2% Gelatin in sterile water (G1393, Sigma-Aldrich). To form a monolayer, fibroblasts were maintained in culture during 24 hours in DMEM supplemented with 10% FBS and 0.01% Penicillin/Streptomycin. HUVECs-CAAX-Venus or HUVECs-CAAX-Venus siRNA-depleted were seeded on top of the fibroblasts confluent monolayer at a density of 15,000 cells/well. Co-cultures were maintained during 24h in EGM-2 medium. When using drugs, cells were treated during 3 hours before analysis. Live-imaging and photoactivation of Cdc42 was performed in a Zeiss 880 confocal laser point scanning microscope using a 63x NA 1.40 oil-immersion objective.

### Drugs Assays

When indicated, cells were treated with (−)-Blebbistatin (Myosin II ATPase inhibitor, 20 µM or 50 µM, Sigma-Aldrich) 10µM of CK-666 (Arp2/3 Complex Inhibitor I, 10µM, Sigma-Aldrich), ML141 (Cdc42 inhibitor, 10µM, Sigma-Aldrich), NSC 23766 (Rac1 inhibitor, 50µM, Tocris) Y-27632 (Rock inhibitor, 20µM, Merck Millipore), Sunitinib malate (Receptor tyrosine kinase inhibitor, 0.05µM Sigma-Aldrich).

### Immunofluorescence

For immunofluorescence of *in vitro* cultured HUVECs, cells were seeded on 24-well plates at concentration of 15,000 cells/well with glass coverslips previously coated with 0.2% Gelatin in sterile water (G1393, Sigma-Aldrich). Both type of cultures, HUVECs alone and co-cultured with fibroblasts were fixed in 4% Paraformaldehyde (PFA) in PBS for 15min at RT. Cells were blocked and permeabilized with blocking solution containing 3% BSA in PBS-T (PBS with 0.1% Triton X-100) for 30min at RT. Then cells were incubated for 2h at RT with the primary antibodies diluted in the blocking solution and washed 3 times for 15min in PBS-T. Primary antibodies used were: anti-activated ITGB1 (BD Pharmingen, 553715, 1:100) and anti-pPaxY118 (Cell Signaling, 2541S, 1:100). Afterwards, cells were incubated in blocking solution containing the secondary fluorophore conjugated antibodies for 1h at RT in the dark, followed again by 3 washes of 15min in PBS-T. Finally, HUVECs were incubated with 1x DAPI (D1306, Molecular Probes by Life Technologies) diluted in PBS-T for 5min in the dark. Coverslips were then mounted on microscopy glass slides using Mowiol DABCO (Sigma-Aldrich). High-resolution Z-stack images at multiple positions were acquired on a confocal laser point-scanning microscope 880 (Zeiss) equipped with the Zen black software with a Plan Apochromat 63x NA 1.40 oil DIC M27 objective. To analyse immunofluorescence intensities, high-resolution Z-stack confocal images of HUVECs stained for pPaxilin, activated ITGB1 and CAAX-Venus were analyzed in Bitplane Imaris using the Mask option to select CAAX-Venus positive cells of interest and quantify levels of fluorescence in masked volumes.

### Rac1 activation assay

For Rac1 activation assay, a commercial kit was used following the manufacturer’s instructions (027BK034-S, Cytoskeleton). Briefly, HUVECs were plated in 100 mm dishes, previously coated with fibronectin (10µg/ml, Sigma-Aldrich), until they reach the confluence. Cells were then starved overnight in serum-free medium, treated with (−)-Blebbistatin (50µM, Sigma-Aldrich) during 2 hours and stimulated with human recombinant VEGF165 (25ng/ml, R&D System) during 30 minutes.

Afterwards, cells were lysate with the lysis buffer provided by the manufactured. Protein concentration was determined by the BCA protein assay kit (Pierce), and 70 μg of lysates were kept for total Rac1 quantification. The remaining sample was incubated, during 1hour at 4°C, with 20 μg of GST-PAK PBD beads, that specifically bind to the active GTP bound form of Rac1 (48). Beads were later washed and the total and activated Rac1 were detected by western blotting (BD Biosciences).

### Protein extraction and Western Blotting

Protein extraction was performed from HUVECs seeded on 6-well plates, which were lysed in 120µL of RIPA buffer supplemented with phosphatase and proteinase inhibitors cocktail (1:100, #10085973 Fischer Scientific). Adherent cells were then detached from the plate with a cell scrapper and the cell lysates were gathered and transferred into an ice cold eppendorf tube. The cell lysates were then centrifuged at maximum speed for 10min at 4°C and the supernatants collected into a new eppendorf tube. Protein concentration was quantified using the BCA protein assay kit (Pierce) following the guidelines recommended by the manufacturer. The Multimode microplate reader, Infinite M200 (Tecan), was used for spectrophotometric measurement of protein with the i-control™ software. For Western Blotting protein samples were normalized up to 25µL and combined with a mixture of 4x Laemmli Sample Buffer (#161-0747, Bio-rad Laboratories) with 450mM DTT (D0632, Sigma-Aldrich) and incubated at 95°C in a Dry Block Thermostat (Grant Instruments, Ltd) for 5min. Protein samples were loaded and separated on a 4-15% Mini-PROTEAN-TGX Gel (#456-1084, BioRad) along with 5µL of protein ladder (Full-Range RPN800E, GE Healthcare Rainbow Molecular Weight Markers). After transfer, blotted membranes were incubated in Ponceau Red to assess transfer quality, and then washed in TBS-T (50mM Tris/HCl, 150mM NaCl, 0.1% Tween-20, pH7.5). Then membranes were incubated in blocking buffer containing 3% BSA (Bovine Serum Albumin, MB04602, Nzytech) in TBS-T for 1h at RT, followed by an overnight incubation at 4°C with the primary antibodies diluted in the same blocking buffer, anti-ACTR3 (Abcam, ab181164, 1:1000), anti-MIIA (Sigma, M8064, 1:1000), anti-GAPDH (Sigma, G8795, 1:1000), anti-βPix (Merck, 07-1450-I, <u>1:1000</u>), anti-Rac1 (BD Bioscience, 610651, 1:1000) and anti-α-tubulin (Sigma, T6199, 1:1000). On the following day membranes were washed 3 times in TBS-T and incubated in blocking buffer containing the secondary horseradish peroxidase (HRP)-conjugated antibodies for 1h at RT. Before revelation membranes were washed again 3 times in TBS-T for 5min and then incubated in ECL™ Western Blotting Detection Reagent (RPN2209, GE Healthcare) following the manufacturer’s protocol. Protein bands were visualized in Chemidoc XRS+ and relative protein quantities were measured using the Image Lab software, both from Bio-Rad Laboratories.

### Statistical analysis

All statistical analysis was performed using GraphPad Prism 7. Measurements were taken from distinct samples, and statistical details of experiments are reported in the figures and figure legends. Sample size is reported in the figure legends and no statistical test was used to determine sample size. The biological replicate is defined as the number of cells, images, animals, as stated in the figure legends. No inclusion/exclusion or randomization criteria were used and all analyzed samples are included. Comparisons between two experimental groups were analyzed with two-sided unpaired parametric t-test or Mann-whitney test, while multiple comparisons between more than two experimental groups were assessed with one-way ANOVA. We considered a result significant when p<0.05. For all box plots: centre line, median; +, mean; whiskers, min to max.

## Author Contributions

A.F. and P.B. contributed equally to this work. A.F., P.B., R.R.F., A.R., S.V., D.R., A.P., A.L., Y.C., F.V. and C.A.F. designed the research, performed experiments, and analysed data. F.E.M., A.M.L.D., and D.V. provided materials and animals. C.A.F. provided funding acquisition, project administration, and resources; C.A.F. wrote the manuscript. A.F., P.B., R.R.F., F.F.V., A.M.L.D., D.V. and C.A.F reviewed and edited the manuscript.

## Acknowledgements

We would like to thank Clare M. Waterman, Robert Fischer, Robert Adelstein and Xuefei Ma (NHLBI) for discussion and technical support; João Barata, Edgar Gomes (IMM) and Michael Potente (MPI-HLR) for critical review of the manuscript. We thank all members of the VML lab for helpful discussions and critical reading of the manuscript. C.A.F was supported by European Research Council starting grant (679368), the European Union H2020-TWINN-2015 – Twinning (692322), the Fundação para a Ciência e a Tecnologia funding (grants: IF/00412/2012; EXPL/BEX-BCM/2258/2013; PRECISE-LISBOA-01-0145-FEDER-016394; PTDC/MED-PAT/31639/2017; PTDC/BIA-CEL/32180/2017; CEECIND/04251/2017); and a grant from the Fondation Leducq (17CVD03).

## SUPPLEMENTARY FIGURE LEGENDS

**Supplementary Figure 1:**
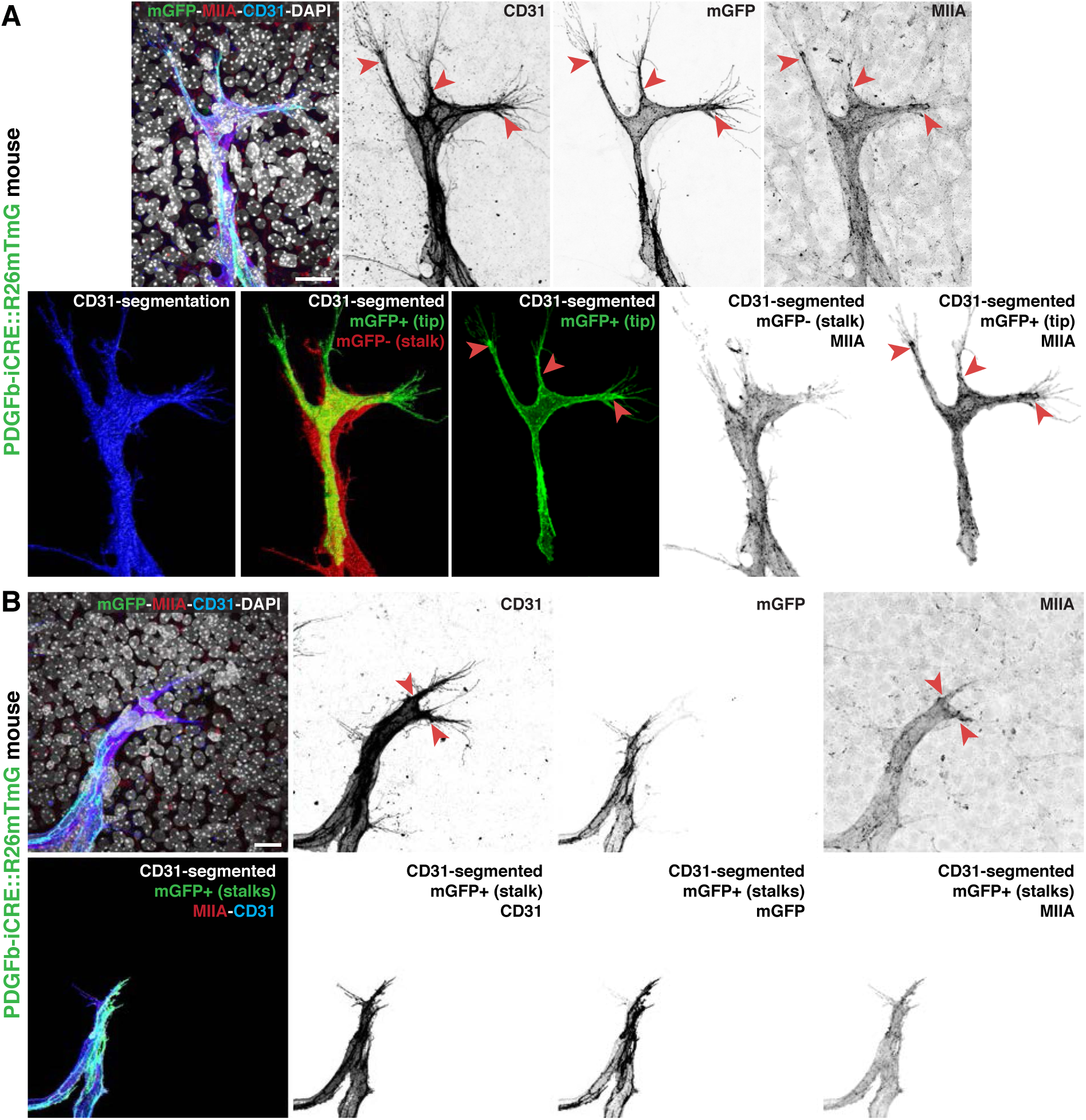
(A) Representative images of a tip cell from *PDGFb-iCre::R26mTmG* mouse retina labeled for MIIA (red), cell membrane (green), CD31 (blue) and DAPI (grey). CD31 segmentation applied to the image and representative images of mGFP^+^ (tip cell) or mGFP^-^ (stalk cell), used for quantification of Figure 1B. Red arrowheads point to sites enriched in MIIA at the base of filopodia. Scale bar, 20 µm. (B) Representative images of 2 stalk cells from *PDGFb-iCre::R26mTmG* mouse retina labeled for MIIA (red), cell membrane (green), CD31 (blue) and DAPI (grey). CD31 segmentation applied to the image and representative images of mGFP^+^ (stalk cells), used for quantification of Figure 1B. Red arrowheads point to sites enriched in MIIA at the base of filopodia. Scale bar, 20 µm.

**Supplementary Figure 2:**
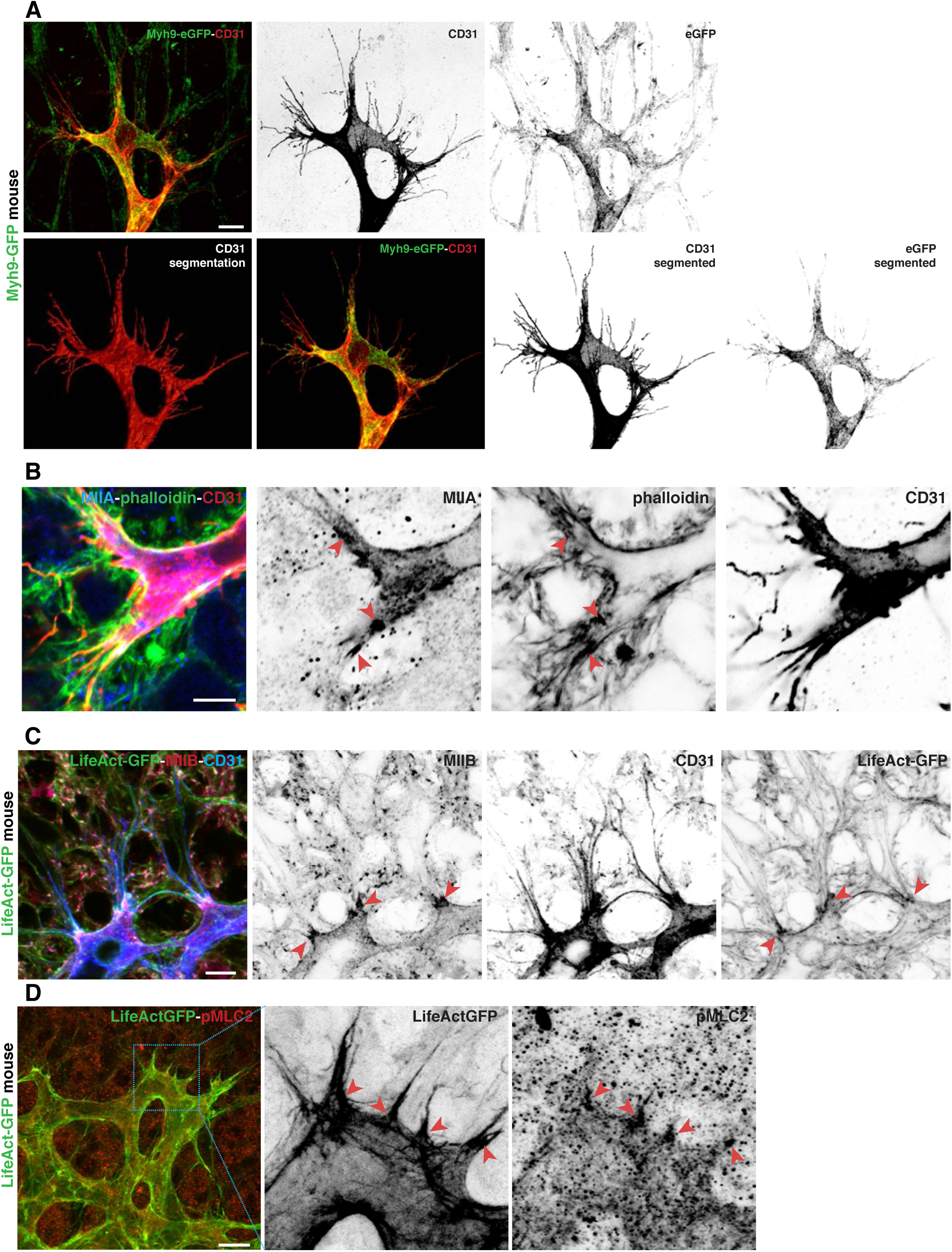
(A) Representative images of a tip cell from *Myh9-GFP* mouse retina labelled for CD31 (red). CD31 segmentation in order to show MIIA localization in the tip cells. Scale bar, 20 µm. (B) High-magnification single z-plane of a tip cell labeled for CD31 (red), F-actin (green) and MIIA (blue) from a WT mouse retina. Red arrowheads point to sites enriched in MIIA at the base of filopodia. Scale bar, 5 µm. (C) Representative images of tip cells in the sprouting front of *LifeAct-GFP* mouse retinas labeled for MIIB (red), actin (green) and CD31 (blue). Red arrowheads point to sites enriched in MIIB at the base of filopodia. Scale bar, 20 µm. (D) Representative images of tip cells in the sprouting front of *LifeAct-GFP* mouse retinas labeled for pMLC2 (red). Red arrowheads point to sites enriched in pMLC2 at the base of filopodia. Scale bar, 50 µm.

**Supplementary Figure 3:**
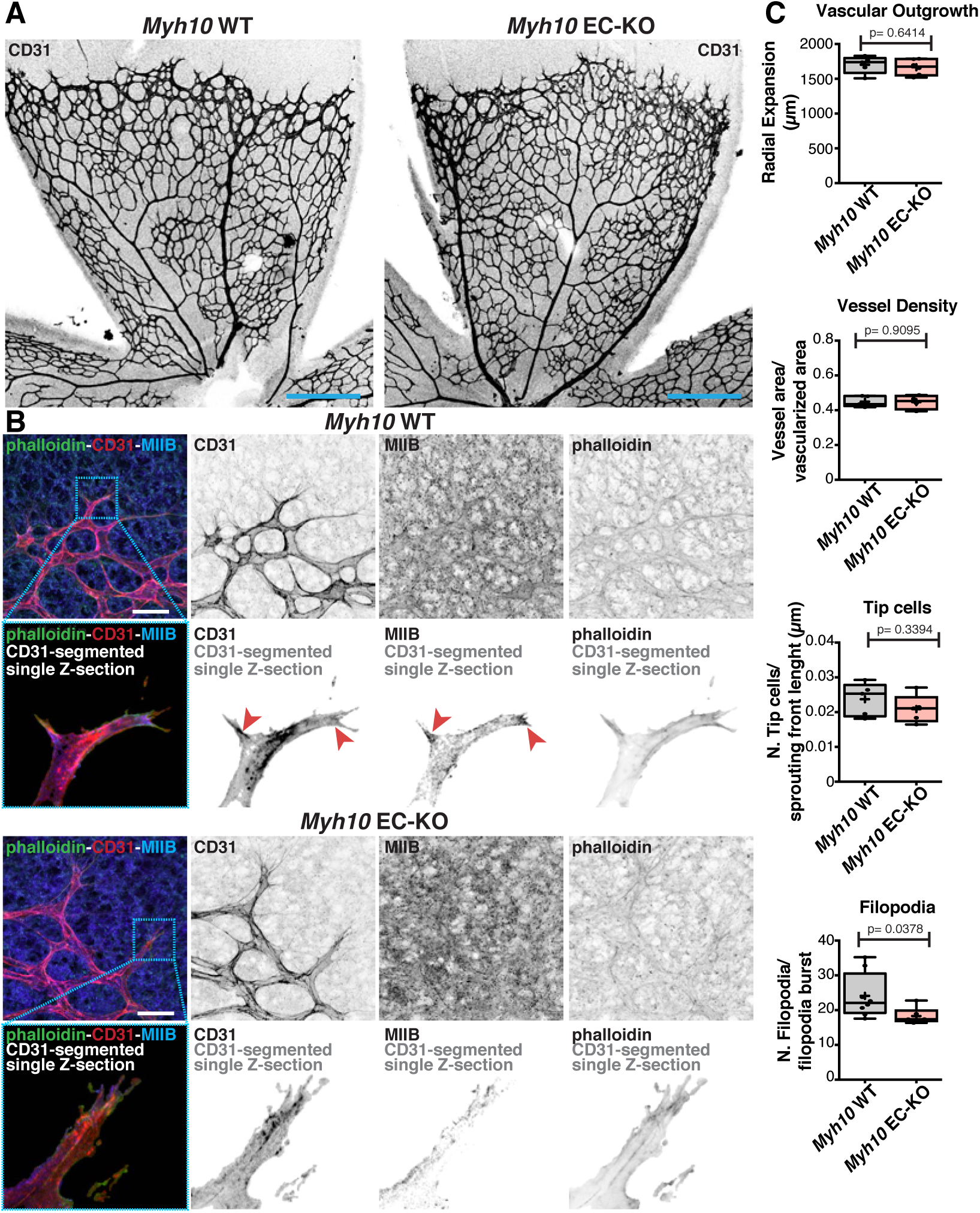
(A) Representative images of mouse retinas from *Myh10* WT and *Myh10* EC-KO labeled for CD31. Scale bar, 250 µm. (B) Higher-magnification and single-plane images of tip cells labeled for CD31 (red), F-actin (green) and MIIB (blue) from *Myh10* WT and *Myh10* EC-KO, from corresponding cyan inserts. Red arrowheads point to sites enriched in MIIB at the base of filopodia. Note a very strong absence of MIIB staining in *Myh10* EC-KO retinas. Scale bar, 20 µm. (C) Box plot of vascular outgrowth, vessel density, number of tip cells and number of filopodia in *Myh10* WT (n = 5 retinas) and *Myh10* EC-KO (n = 6 retinas) mouse retinas. *p*-values from unpaired Student t-test.

**Supplementary Figure 4:**
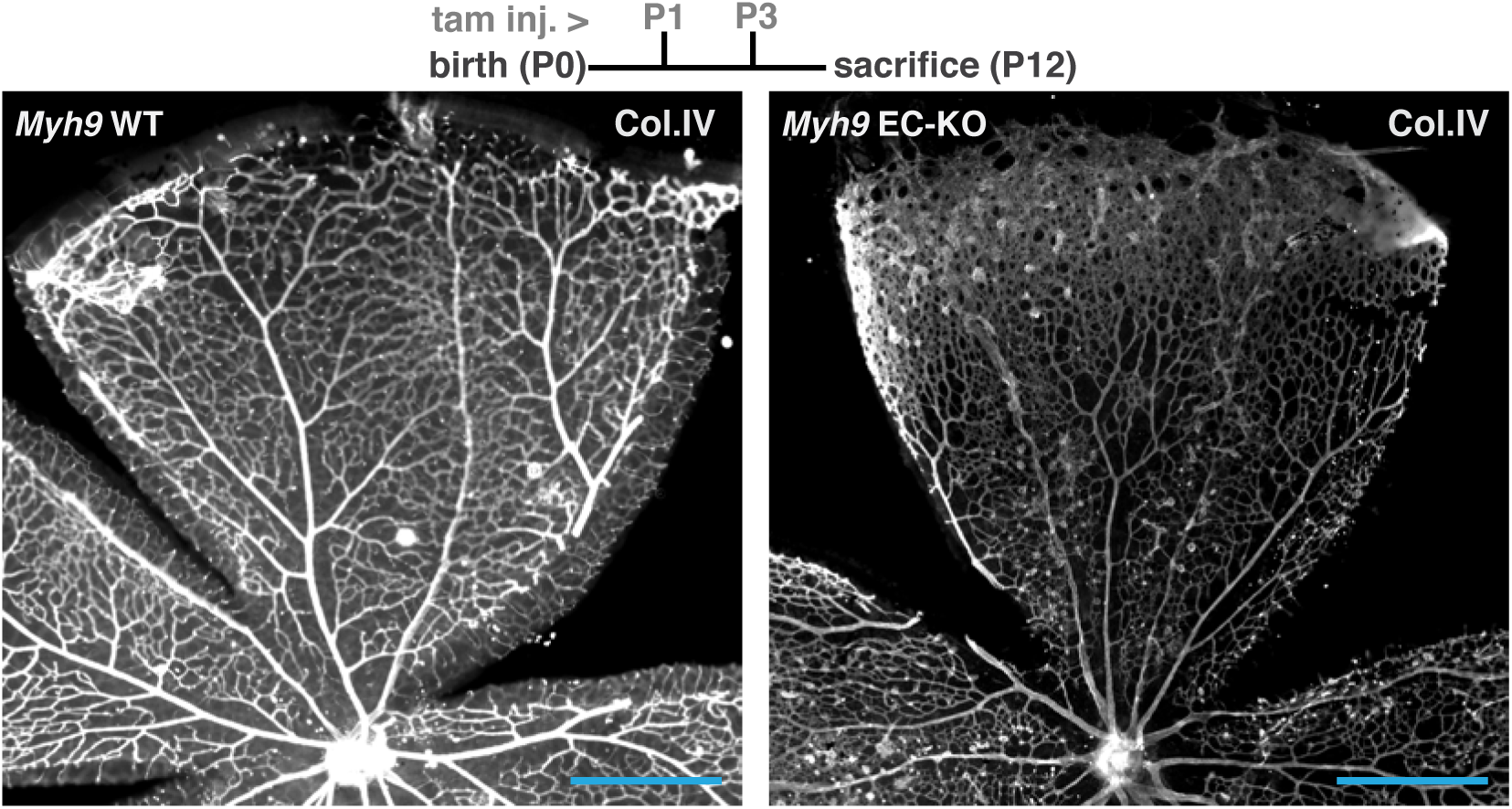
(A) Representative images of tip cells from *Myh9* WT and *Myh9* EC-KO mouse retinas labeled for CD31 and MIIA. Scale bar, 5 µm. (B) Representative images of vascular plexus from *Myh9* WT and *Myh9* EC-KO mouse retinas labeled for CD31, red arrowheads highlight filopodia. Scale bar, 50 µm. (C) Box plot of number of plexus filopodia per area (normalized to control) in *Myh9* WT (n = 4 retinas) and *Myh9* EC-KO (n = 4 retinas) mouse retinas. *p*-values from unpaired Student t-test. (D) Representative images of tip cells in the sprouting front of *Myh9* EC-KO labelled for CD31 (red) and MIIB (green). Scale bar, 20 µm.

**Supplementary Figure 5:**
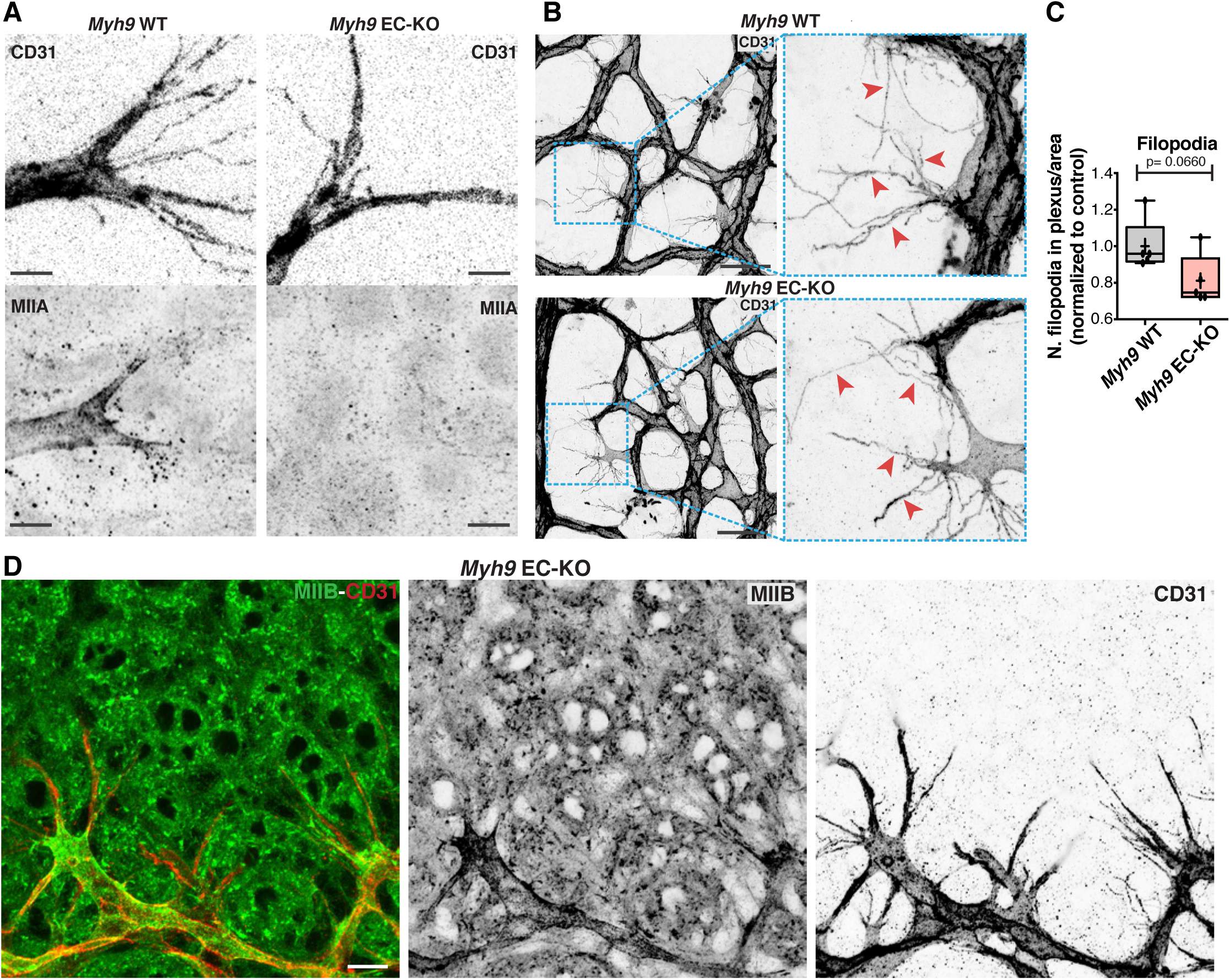
(A) Upper panel: timeline of tamoxifen injection (tam. inj.) and age of sacrifice of mouse pups. Representative images of overview of P12 retinas from *Myh9* WT and *Myh9* EC-KO labeled for Col.IV. Scale bar, 500 µm.

**Supplementary Figure 6:**
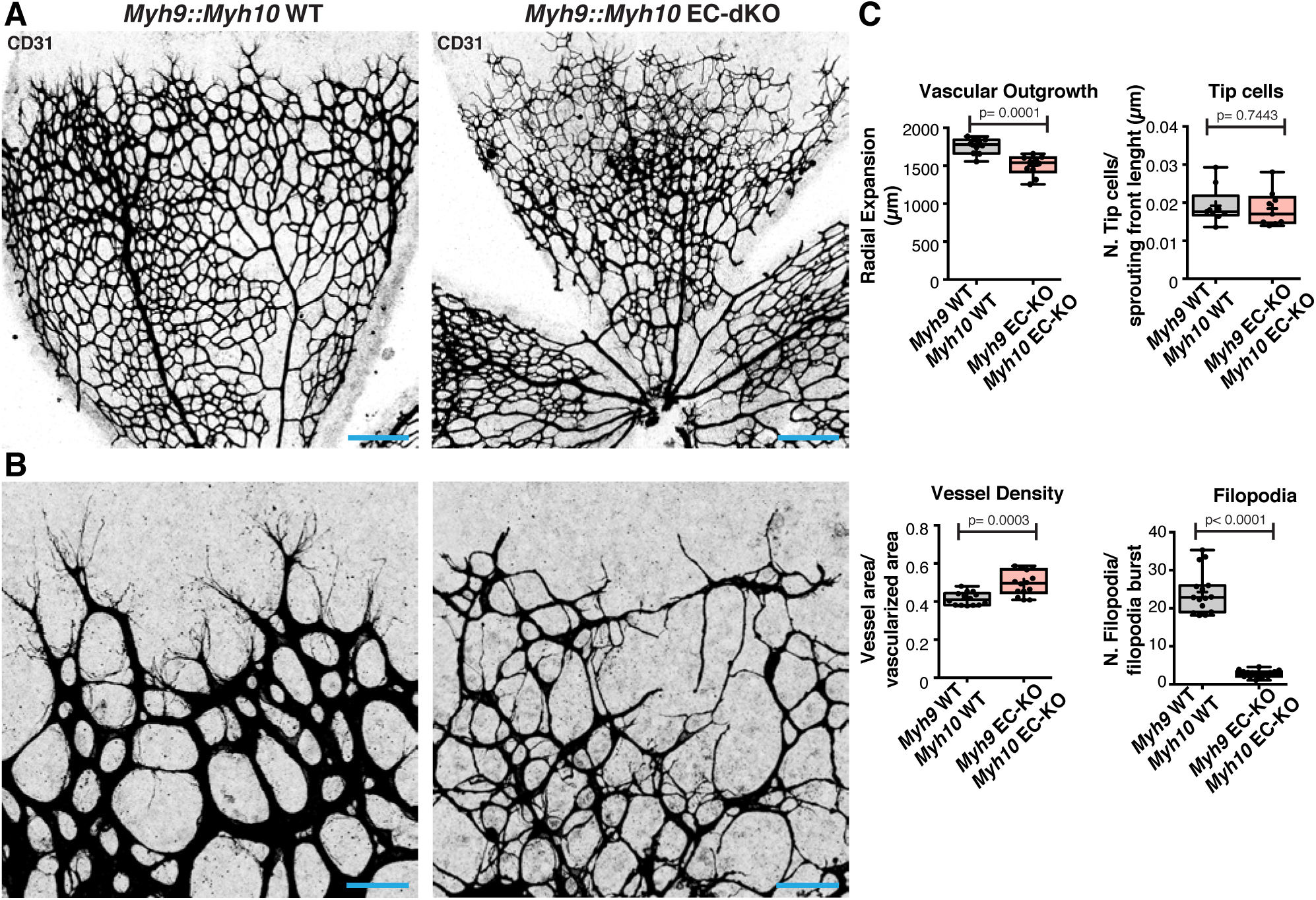
(A) Representative images of mouse retinas from *Myh9::Myh10* WT and *Myh9::Myh10* EC-dKO labeled for CD31. Scale bar, 250 µm. (B) Representative images of tip cells from sprouting fronts of *Myh9::Myh10* WT and *Myh9::Myh10* EC-dKO labeled for CD31. Scale bar, 50 µm. (C) Box plot of vascular outgrowth, vessel density (vessel area per vascularized area), number of tip cells per sprouting front length (µm) and number of filopodia per filopodia burst in *Myh9::Myh10* WT (n = 9-15 retinas) and *Myh9::Myh10* EC-dKO (n = 10-17 retinas) mouse retinas. *p*-values from unpaired Student t-test.

**Supplementary Figure 7:**
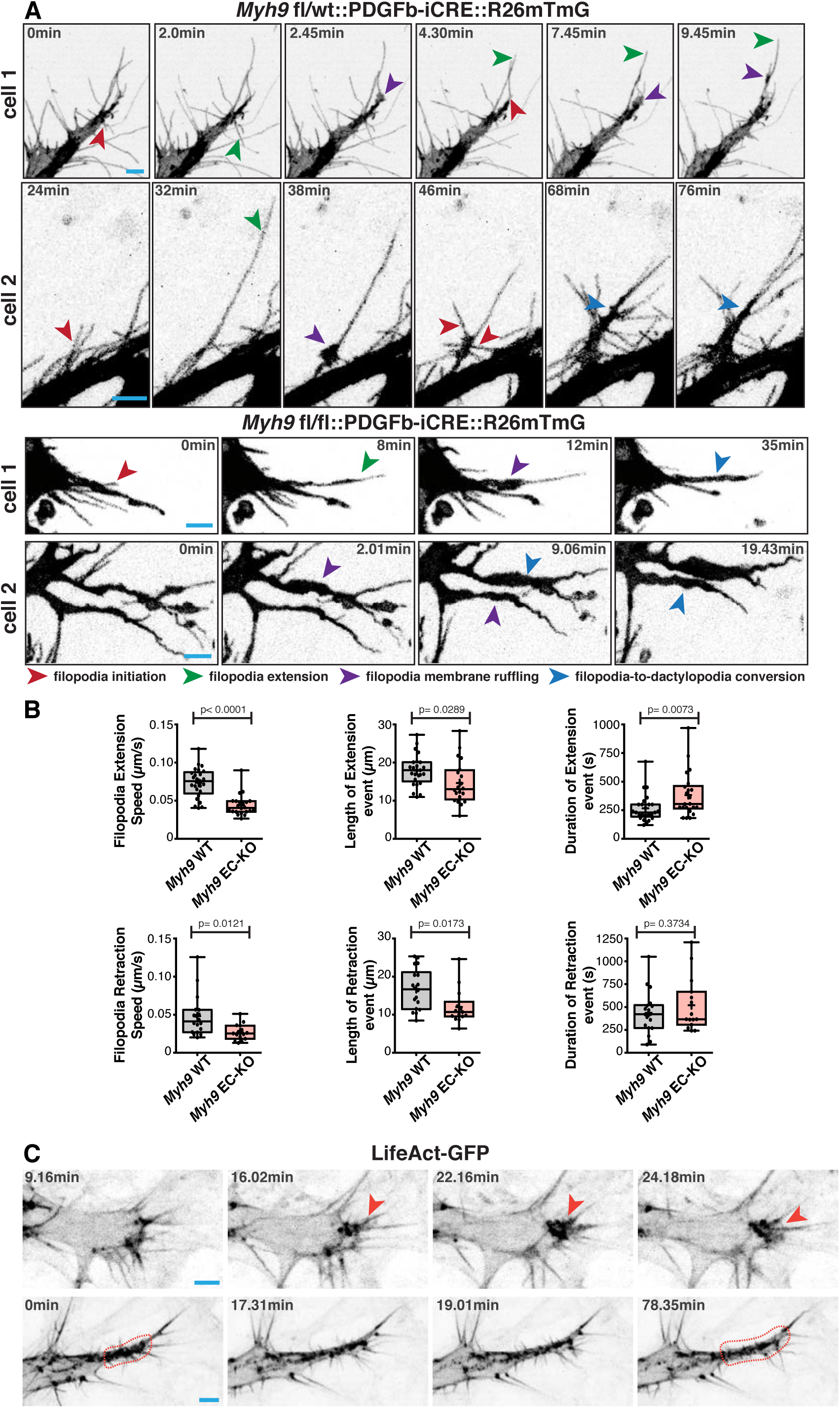
(A) Time-course representative images of LLPs and filopodia dynamics in *Myh9 fl/wt::PDGFb-iCre::R26mTmG* and *Myh9 fl/fl::PDGFb-iCre::R26mTmG* EC-KO mouse retinas. Red arrows indicates filopodia initiation; Green arrows indicates filopodia extension; Purple arrows indicates filopodia membrane ruffling; and Blue arrows indicates filopodia-to-LLPs conversion. Scale bar, 5 µm. (B) Box plots of filopodia extension speed (µm/s), length of extension event (µm), duration of extension event (s), filopodia retraction speed (µm/s), length of retraction event (µm) and duration of retraction event (s) in *Myh9 fl/wt::PDGFb-iCre::R26mTmG* and *Myh9 fl/fl::PDGFb-iCre::R26mTmG* EC-KO mouse retinas. *Myh9* WT n = 18 filopodia; *Myh9* EC-KO n = 15 filopodia. *p*-values from unpaired Student t-test. (C) Representative images of actin dynamics in the formation of filopodia and LLPs in *LifeAct-GFP* mouse tip cells. Red arrow indicates filopodia initiation and red dashed line contour represents LLPs. Scale bar, 5 µm.

**Supplementary Figure 8:**
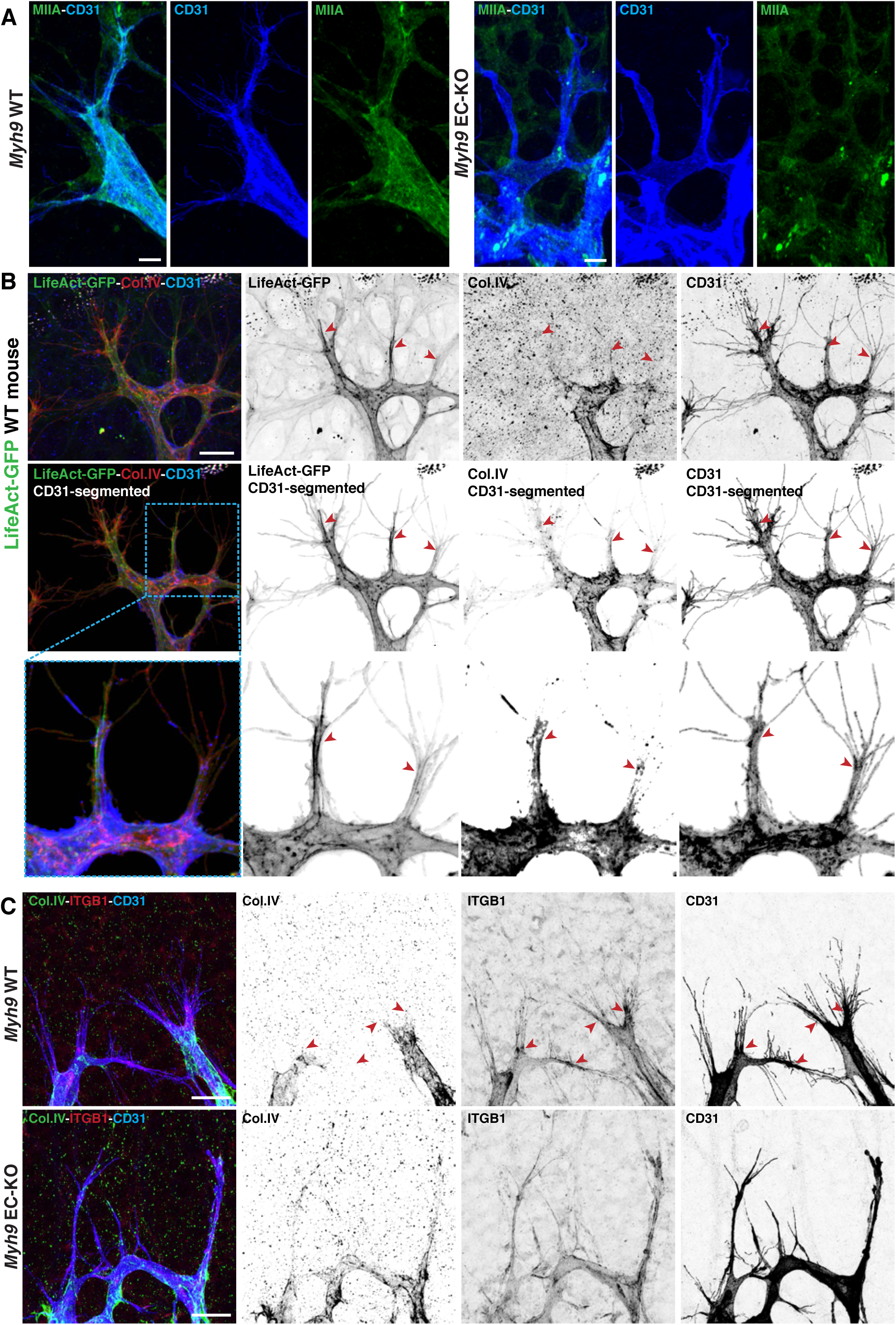
(A) Representative images of tip cells of mouse retinas from *Myh9* WT and *Myh9* EC-KO labeled for MIIA (green) and CD31 (blue) and correspondent CD31-segmented image to see MIIA localization in tip cells, used to highlight LLPs in Figure 2A. Scale bar, 10 µm. (B) Representative images of a tip cell from *LifeAct-GFP Myh9* WT mouse retina labeled for Col.IV (red), actin (green) and CD31 (blue) and correspondent CD31-segmented image. Red arrowheads point to LLPs in contact with non-vascular ECM. Scale bar, 20 µm. (C) Representative images of tip cells from mouse retinas from *Myh9* WT and *Myh9* EC-KO labeled for Col.IV (green), ITGB1 (red) and CD31 (blue). Red arrowheads point to sites enriched in aITGB1 in contact with non-vascular ECM at the base of filopodia in *Myh9* WT tip cells, which is absent and more homogenously distributed along LLPs of *Myh9* EC-KO tip cells. Scale bar, 20 µm.

**Supplementary Figure 9:**
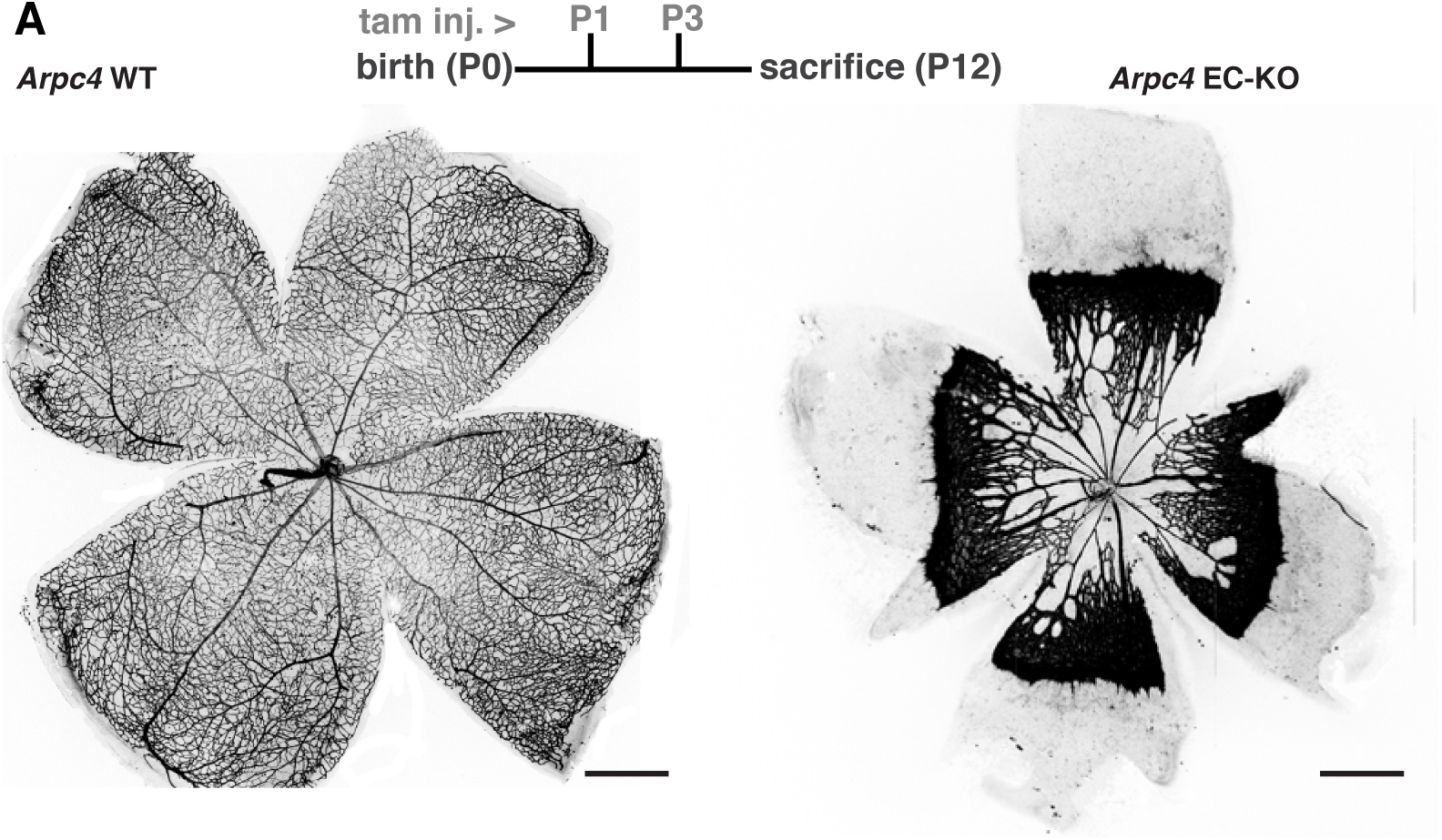
(A) Upper panel: timeline of tamoxifen injection (tam. inj.) and age of sacrifice of mouse pups. Representative images of mouse retinas from *Arpc4* WT and *Arpc4* EC-KO labeled for CD31 at P12. Scale bar, 500 µm.

**Supplementary Figure 10:**
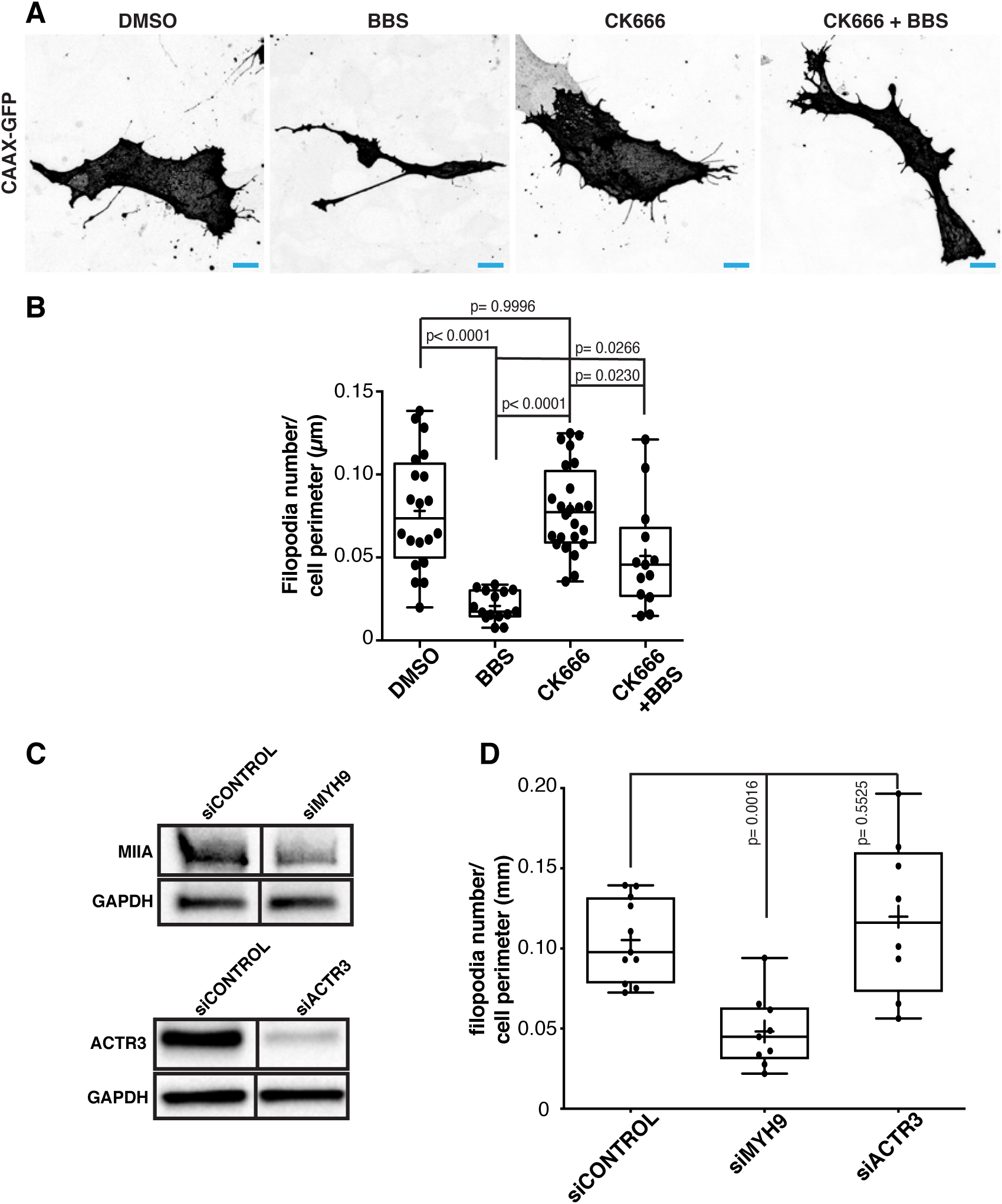
(A) Representative images of HUVECs transduced with lenti-CAAX-Venus co-cultured with fibroblasts treated with in DMSO, BBS, CK666, and BBS + CK666 conditions. Scale bar, 10 µm. (B) Box plot of filopodia number in DMSO (n = 20 cells), BBS (n = 15 cells), CK666 (n = 24 cells), and BBS + CK666 (n = 13 cells). *p*-values from unpaired ANOVA test.

**Supplementary Figure 11:**
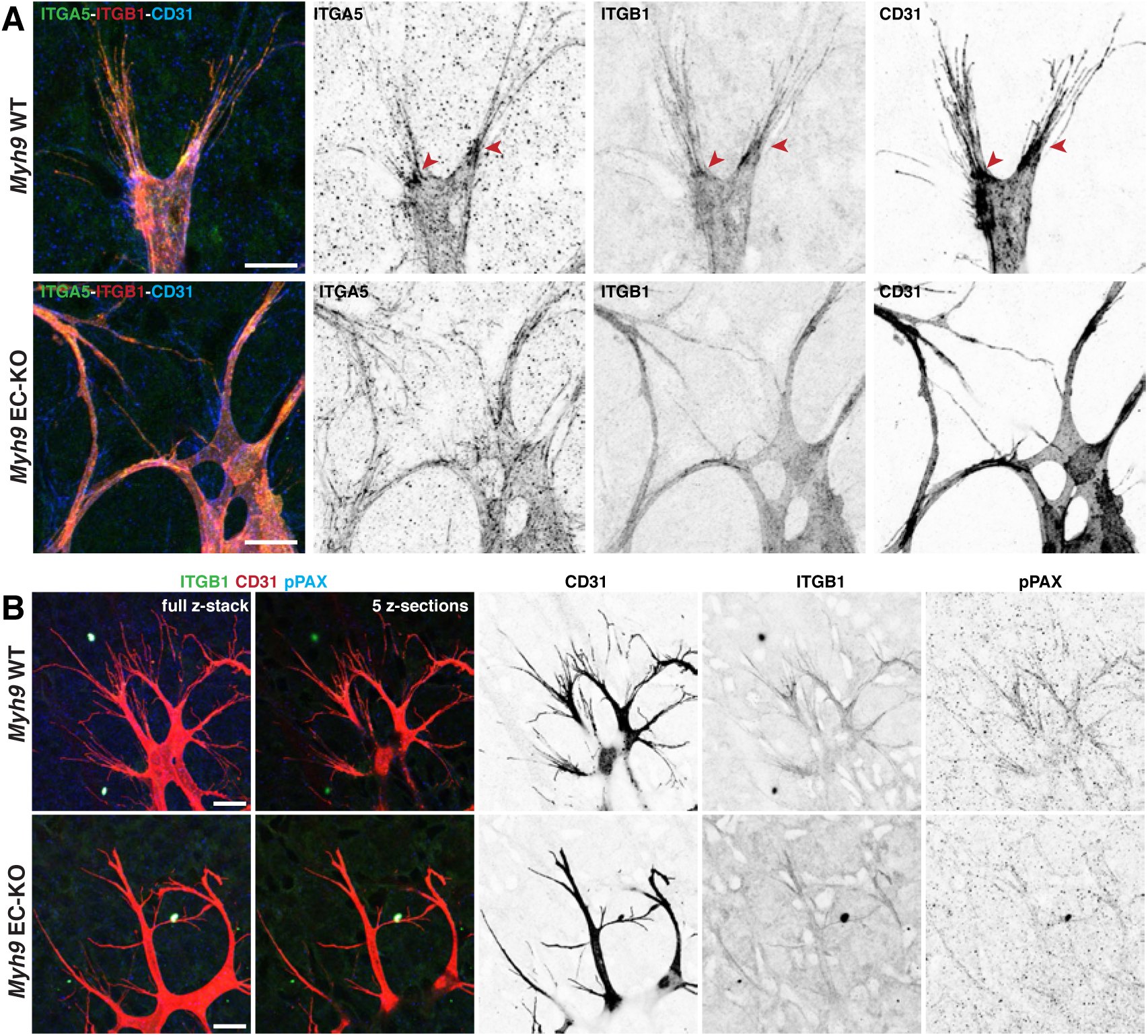
(A) Representative images of tip cells from sprouting fronts of *Myh9* WT and *Myh9* EC-KO mouse retinas labeled for ITGA5 (green), activated ITGB1 (red) and CD31 (blue). Red arrowheads point to sites enriched in aITGB1 and ITGA5 at the base of filopodia in *Myh9* WT tip cells, which is absent or more homogenously distributed along LLPs of *Myh9* EC-KO tip cells. Scale bar, 20 µm. (B) Representative images (full z-stack and single z-section) of tip cells from sprouting fronts of *Myh9* WT and *Myh9* EC-KO mouse retinas labeled for activated ITGB1 (green), CD31 (red) and pPax Y118 (blue) and also 5 z-sections correspondent images, which were used to highlight integrin and pPAX stainings in Figure 4A. Scale bar, 20 µm.

**Supplementary Figure 12:**
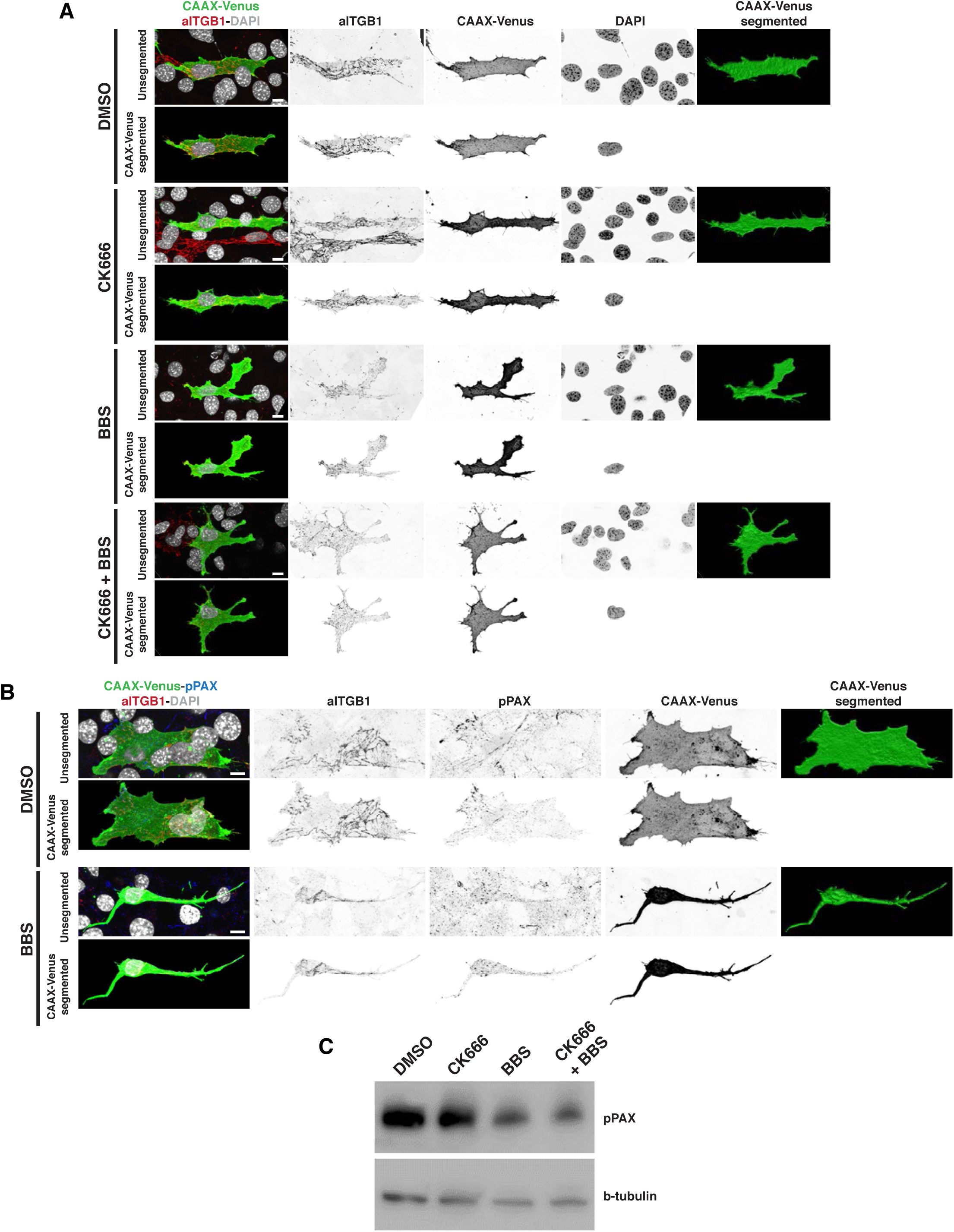
(A) Representative images of HUVECs expressing CAAX-Venus at the cell membrane in DMSO, BBS, CK-666 and CK-666+BBS conditions. HUVECs labeled for activated ITGB1 (red), cell membrane (CAAX-Venus, in green) and nuclei (DAPI, in grey). Correspondent Venus-segmented images for a better analysis, which were used to highlight individual cells displayed in Figure 4C. Scale bar, 10 µm. (B) Representative images of HUVECs expressing CAAX-Venus at the cell membrane in DMSO and BBS conditions. HUVECs labeled for activated ITGB1 (red), pPaxilin (pPAX, blue) cell membrane (CAAX-Venus, in green) and nuclei (DAPI, in grey). Correspondent Venus-segmented images for a better analysis. Scale bar, 10 µm. (C) Western-blot of pPAX levels in HUVEC cultures treated for 3h in DMSO, BBS, CK-666 or CK-666+BBS conditions.

**Supplementary Figure 13:**
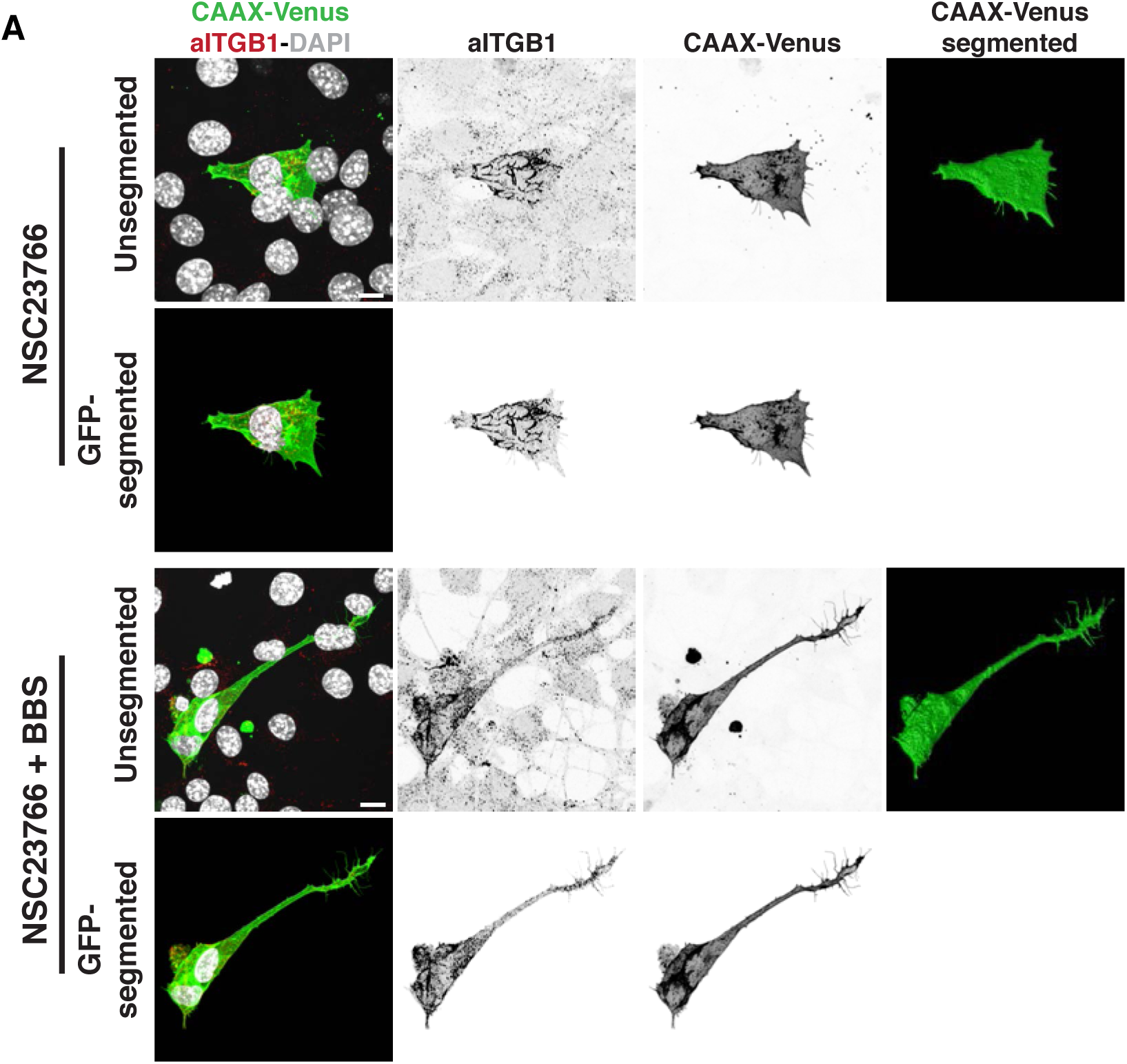
(A) Representative images of HUVECs expressing CAAX-Venus at the cell membrane in NSC23766 and NSC23766 + BBS conditions. HUVECs labeled for activated ITGB1 (red), cell membrane (CAAX-Venus, in green) and nuclei (DAPI, in grey). Correspondent Venus-segmented images for a better analysis, which were used to highlight individual cells displayed in Figure 4F. Scale bar, 10 µm.

**Supplementary Figure 14:**
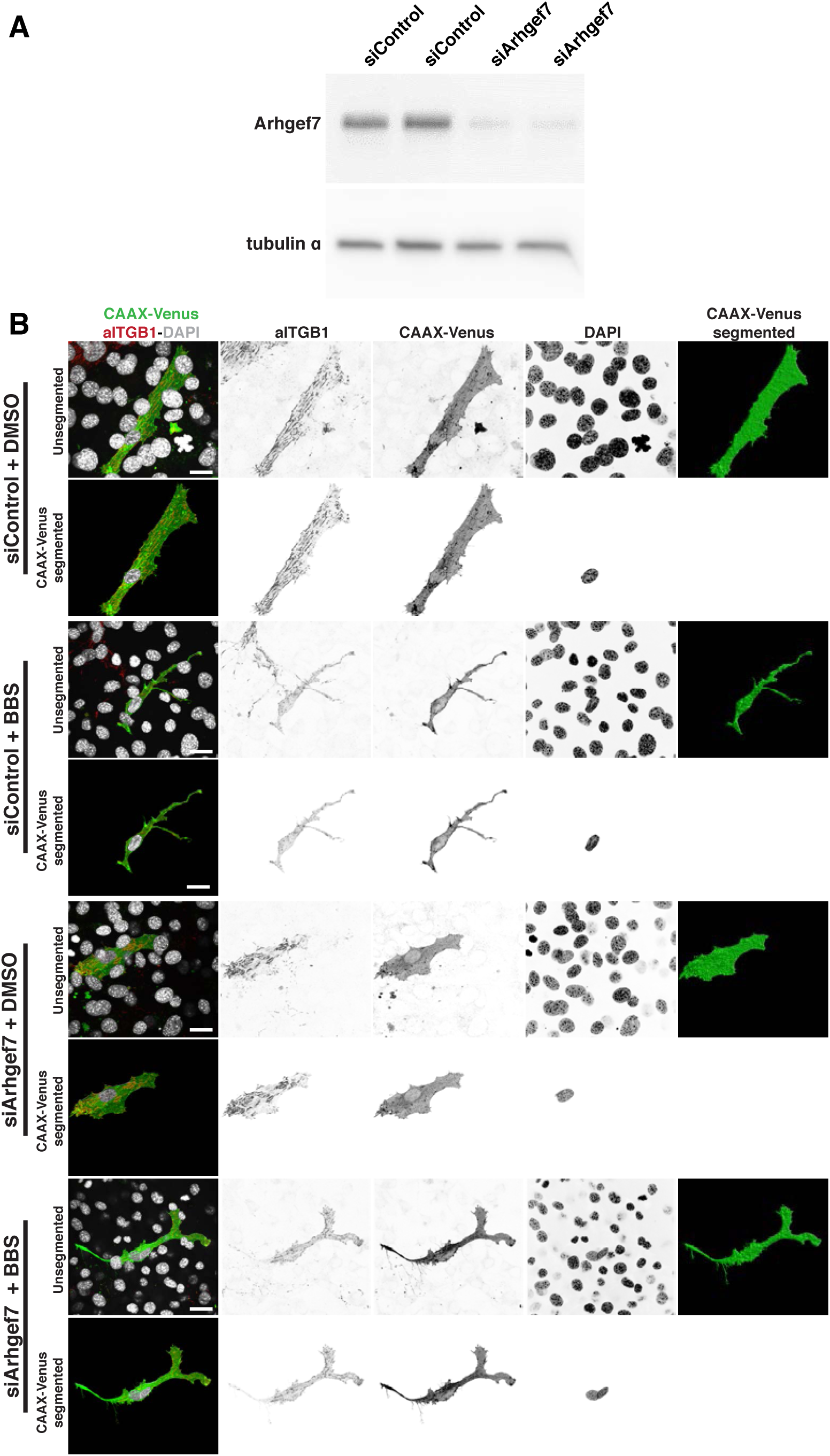
(A) Western blot analysis of efficiency of knock-down for Arhgef7 (bPIX) in siControl and siArhgef7-treated HUVECs. Tubulin was used as loading control. (B) Representative images of HUVECs expressing CAAX-Venus at the cell membrane in siControl and siArhgef7-treated HUVECs treated with DMSO or BBS. HUVECs labeled for activated ITGB1 (red), cell membrane (CAAX-Venus, in green) and nuclei (DAPI, in grey). Correspondent Venus-segmented images for a better analysis, which were used to highlight individual cells displayed in Figure 4H. Scale bar, 20 µm.

**Supplementary Figure 15:**
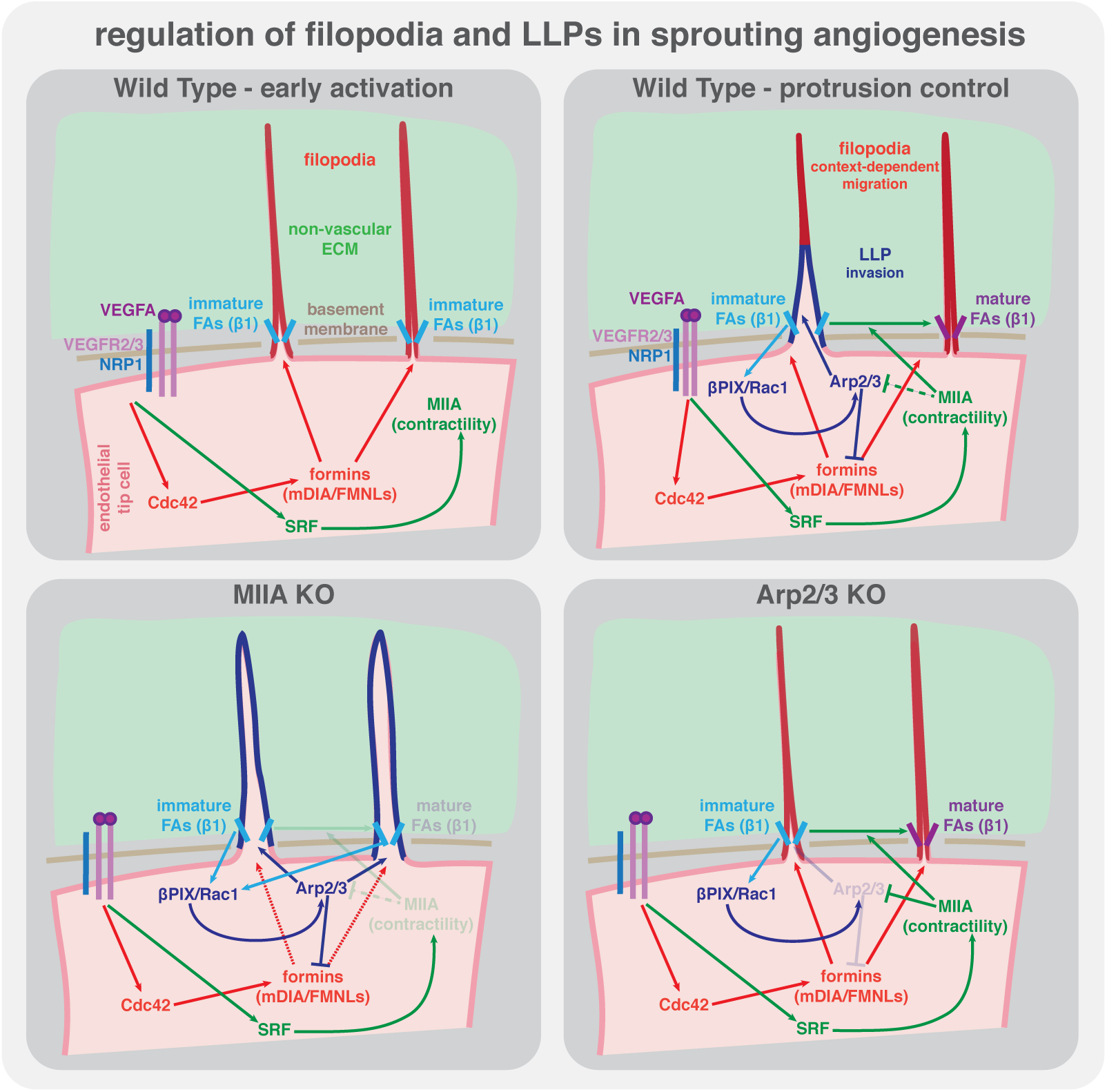
Working model for the fine-tuning between filopodia and LLPs in endothelial tip cells. MIIA regulates maturation of focal adhesions at the base of filopodia, which regulates negatively Arp2/3 activity downstream of bPix/Rac1 signalling pathway, preventing LLPs formation and invasion. In the scenario where MIIA is not present, there is an overactivation of Arp2/3 and unregulated conversion of filopodia into LLPs. When Arp2/3 is absent, there is an abrogation of LLPs formation an inhibition of migration.

## SUPPLEMENTARY MOVIES

**Supplementary Video 1:** 3D rotational view of a tip cell from LifeAct-GFP mouse retinas labeled for MIIA (red), actin (green) and CD31 (blue).

**Supplementary Video 2:** 3D rotational view of a tip cell from Cdh5-iCre::R26mTmG mouse retina labeled for MIIA (red), cell membrane (green), CD31 (blue) and DAPI (grey).

**Supplementary Video 3:** Live-imaging of an endothelial tip cell in *Myh9 fl/wt::PDGFB-iCRE::R26mTmG* P6 mouse retina, showing filopodia and LLPs dynamics.

**Supplementary Video 4:** Live-imaging of an endothelial tip cell in *Myh9 fl/wt::PDGFB-iCRE::R26mTmG* P6 mouse retina, showing filopodia membrane ruffling.

**Supplementary Video 5:** Live-imaging of an endothelial tip cell in *Myh9 fl/wt::PDGFB-iCRE::R26mTmG* P6 mouse retina, showing emergence of LLPs from filopodia.

**Supplementary Video 6:** Live-imaging of an endothelial tip cell in *Myh9 fl/wt::PDGFB-iCRE::R26mTmG* P6 mouse retina, showing filopodia membrane ruffling, filopodia and LLPs dynamics.

**Supplementary Video 7:** Live-imaging of an endothelial tip cell in *Myh9 fl/fl::PDGFB-iCRE::R26mTmG* P6 mouse retina, showing filopodia and LLPs dynamics.

**Supplementary Video 8:** Live-imaging of an endothelial tip cell in *Myh9 fl/fl::PDGFB-iCRE::R26mTmG* P6 mouse retina, showing abnormal filopodia membrane ruffling, and filopodia to LLPs conversion, example 1.

**Supplementary Video 9:** Live-imaging of an endothelial tip cell in *Myh9 fl/fl::PDGFB-iCRE::R26mTmG* P6 mouse retina, showing abnormal filopodia membrane ruffling, and filopodia to LLPs conversion, example 2.

**Supplementary Video 10:** Live-imaging of an endothelial tip cell in *Myh9 fl/fl::PDGFB-iCRE::R26mTmG* P6 mouse retina, showing abnormal LLPs dynamics.

**Supplementary Video 11:** Live-imaging of an endothelial tip cell in *Myh9 fl/fl::PDGFB-iCRE::R26mTmG* P6 mouse retina, showing abnormal filopodia membrane ruffling, and filopodia to LLPs conversion, example 3.

**Supplementary Video 12:** Live-imaging of actin dynamics in retinal endothelial tip cells in a *LifeAct-GFP* P6 mouse retina, example 1.

**Supplementary Video 13:** Live-imaging of actin dynamics in retinal endothelial tip cells in a *LifeAct-GFP* P6 mouse retina, example 2.

**Supplementary Video 14:** Live-imaging of filopodia and lamellipodia dynamics in a HUVEC expressing an optogenetic activator of Cdc42 (ILID) in DMSO.

**Supplementary Video 15:** Live-imaging of filopodia and lamellipodia dynamics in a HUVEC expressing an optogenetic activator of Cdc42 (ILID) in CK666.

**Supplementary Video 16:** Live-imaging of filopodia and lamellipodia dynamics in a HUVEC expressing an optogenetic activator of Cdc42 (ILID) in BBS.

**Supplementary Video 17:** Live-imaging of filopodia and lamellipodia dynamics in a HUVEC expressing an optogenetic activator of Cdc42 (ILID) in CK666 and BBS.

## References

1. Augustin HG, and Koh GY. Organotypic vasculature: From descriptive heterogeneity to functional pathophysiology. Science. 2017;357(6353).

2. Potente M, and Makinen T. Vascular heterogeneity and specialization in development and disease. Nat Rev Mol Cell Biol. 2017;18(8):477–94.

3. Gerhardt H, Golding M, Fruttiger M, Ruhrberg C, Lundkvist A, Abramsson A, et al. VEGF guides angiogenic sprouting utilizing endothelial tip cell filopodia. J Cell Biol. 2003;161(6):1163–77.

4. Jacquemet G, Hamidi H, and Ivaska J. Filopodia in cell adhesion, 3D migration and cancer cell invasion. Curr Opin Cell Biol. 2015;36:23–31.

5. Pollard TD. Actin and Actin-Binding Proteins. Cold Spring Harb Perspect Biol. 2016;8(8).

6. Rottner K, and Schaks M. Assembling actin filaments for protrusion. Curr Opin Cell Biol. 2019;56:53–63.

7. Yamada KM, and Sixt M. Mechanisms of 3D cell migration. Nat Rev Mol Cell Biol. 2019;20(12):738–52.

8. Mattila PK, and Lappalainen P. Filopodia: molecular architecture and cellular functions. Nat Rev Mol Cell Biol. 2008;9(6):446–54.

9. Elliott H, Fischer RS, Myers KA, Desai RA, Gao L, Chen CS, et al. Myosin II controls cellular branching morphogenesis and migration in three dimensions by minimizing cell-surface curvature. Nat Cell Biol. 2015;17(2):137–47.

10. Fischer RS, Gardel M, Ma X, Adelstein RS, and Waterman CM. Local cortical tension by myosin II guides 3D endothelial cell branching. Curr Biol. 2009;19(3):260–5.

11. Alieva NO, Efremov AK, Hu S, Oh D, Chen Z, Natarajan M, et al. Myosin IIA and formin dependent mechanosensitivity of filopodia adhesion. Nat Commun. 2019;10(1):3593.

12. Fonseca CG, Barbacena P, and Franco CA. Endothelial cells on the move: dynamics in vascular morphogenesis and disease. 2020;2(1):H29.

13. Laviña B, Castro M, Niaudet C, Cruys B, Álvarez-Aznar A, Carmeliet P, et al. Defective endothelial cell migration in the absence of Cdc42 leads to capillary-venous malformations. Development. 2018;145:dev161182.

14. Fantin A, Lampropoulou A, Gestri G, Raimondi C, Senatore V, Zachary I, et al. NRP1 regulates CDC42 activation to promote filopodia formation in endothelial tip cells. Cell Reports. 2015;11:1577–90.

15. Nohata N, Uchida Y, Stratman A, Adams R, Zheng Y, Weinstein B, et al. Temporal-specific roles of Rac1 during vascular development and retinal angiogenesis. Developmental Biology. 2016;411:183–94.

16. Franco CA, Blanc J, Parlakian A, Blanco R, Aspalter IM, Kazakova N, et al. SRF selectively controls tip cell invasive behavior in angiogenesis. Development. 2013;140(11):2321–33.

17. Weinl C, Riehle H, Park D, Stritt C, Beck S, Huber G, et al. Endothelial SRF/MRTF ablation causes vascular disease phenotypes in murine retinae. J Clin Invest. 2013;123(5):2193–206.

18. Franco CA, Mericskay M, Parlakian A, Gary-Bobo G, Gao-Li J, Paulin D, et al. Serum response factor is required for sprouting angiogenesis and vascular integrity. Dev Cell. 2008;15(3):448–61.

19. Park H, Yamamoto H, Mohn L, Ambühl L, Kanai K, Schmidt I, et al. Integrin-linked kinase controls retinal angiogenesis and is linked to Wnt signaling and exudative vitreoretinopathy. Nature Communications. 2019;10:5243.

20. Yamamoto H, Ehling M, Kato K, Kanai K, van Lessen M, Frye M, et al. Integrin β1 controls VE-cadherin localization and blood vessel stability. Nature Communications. 2015;6:6429.

21. Phng LK, Stanchi F, and Gerhardt H. Filopodia are dispensable for endothelial tip cell guidance. Development. 2013;140(19):4031–40.

22. Vanlandewijck M, He L, Mae MA, Andrae J, Ando K, Del Gaudio F, et al. A molecular atlas of cell types and zonation in the brain vasculature. Nature. 2018;554(7693):475–80.

23. Wang A, Ma X, Conti MA, Liu C, Kawamoto S, and Adelstein RS. Nonmuscle myosin II isoform and domain specificity during early mouse development. Proc Natl Acad Sci U S A. 2010;107(33):14645–50.

24. Claxton S, Kostourou V, Jadeja S, Chambon P, Hodivala-Dilke K, and Fruttiger M. Efficient, inducible Cre-recombinase activation in vascular endothelium. Genesis. 2008;46(2):74–80.

25. Even-Ram S, Doyle AD, Conti MA, Matsumoto K, Adelstein RS, and Yamada KM. Myosin IIA regulates cell motility and actomyosin-microtubule crosstalk. Nat Cell Biol. 2007;9(3):299–309.

26. Sandquist JC, and Means AR. The C-terminal tail region of nonmuscle myosin II directs isoform-specific distribution in migrating cells. Mol Biol Cell. 2008;19(12):5156–67.

27. Vicente-Manzanares M, Koach MA, Whitmore L, Lamers ML, and Horwitz AF. Segregation and activation of myosin IIB creates a rear in migrating cells. J Cell Biol. 2008;183(3):543–54.

28. Krndija D, El Marjou F, Guirao B, Richon S, Leroy O, Bellaiche Y, et al. Active cell migration is critical for steady-state epithelial turnover in the gut. Science. 2019;365(6454):705–10.

29. Bishop ET, Bell GT, Bloor S, Broom IJ, Hendry NF, and Wheatley DN. An in vitro model of angiogenesis: basic features. Angiogenesis. 1999;3(4):335–44.

30. Guntas G, Hallett RA, Zimmerman SP, Williams T, Yumerefendi H, Bear JE, et al. Engineering an improved light-induced dimer (iLID) for controlling the localization and activity of signaling proteins. Proc Natl Acad Sci U S A. 2015;112(1):112–7.

31. Lavina B, Castro M, Niaudet C, Cruys B, Alvarez-Aznar A, Carmeliet P, et al. Defective endothelial cell migration in the absence of Cdc42 leads to capillary-venous malformations. Development. 2018;145(13).

32. Hodge RG, and Ridley AJ. Regulating Rho GTPases and their regulators. Nat Rev Mol Cell Biol. 2016;17(8):496–510.

33. Etienne-Manneville S, and Hall A. Integrin-mediated activation of Cdc42 controls cell polarity in migrating astrocytes through PKCzeta. Cell. 2001;106(4):489–98.

34. Yamamoto H, Ehling M, Kato K, Kanai K, van Lessen M, Frye M, et al. Integrin beta1 controls VE-cadherin localization and blood vessel stability. Nat Commun. 2015;6:6429.

35. Chrzanowska-Wodnicka M, and Burridge K. Rho-stimulated contractility drives the formation of stress fibers and focal adhesions. J Cell Biol. 1996;133(6):1403–15.

36. Kuo JC, Han X, Hsiao CT, Yates JR, 3rd, and Waterman CM. Analysis of the myosin-II-responsive focal adhesion proteome reveals a role for beta-Pix in negative regulation of focal adhesion maturation. Nat Cell Biol. 2011;13(4):383–93.

37. Huveneers S, and Danen EH. Adhesion signaling - crosstalk between integrins, Src and Rho. J Cell Sci. 2009;122(Pt 8):1059–69.

38. Rotty JD, Wu C, Haynes EM, Suarez C, Winkelman JD, Johnson HE, et al. Profilin-1 serves as a gatekeeper for actin assembly by Arp2/3-dependent and -independent pathways. Dev Cell. 2015;32(1):54–67.

39. Suarez C, Carroll RT, Burke TA, Christensen JR, Bestul AJ, Sees JA, et al. Profilin regulates F-actin network homeostasis by favoring formin over Arp2/3 complex. Dev Cell. 2015;32(1):43–53.

40. Fantin A, Lampropoulou A, Gestri G, Raimondi C, Senatore V, Zachary I, et al. NRP1 Regulates CDC42 Activation to Promote Filopodia Formation in Endothelial Tip Cells. Cell Rep. 2015;11(10):1577–90.

41. Carlier MF, and Shekhar S. Global treadmilling coordinates actin turnover and controls the size of actin networks. Nat Rev Mol Cell Biol. 2017;18(6):389–401.

42. Leon C, Eckly A, Hechler B, Aleil B, Freund M, Ravanat C, et al. Megakaryocyte-restricted MYH9 inactivation dramatically affects hemostasis while preserving platelet aggregation and secretion. Blood. 2007;110(9):3183–91.

43. Tullio AN, Accili D, Ferrans VJ, Yu ZX, Takeda K, Grinberg A, et al. Nonmuscle myosin II-B is required for normal development of the mouse heart. Proc Natl Acad Sci U S A. 1997;94(23):12407–12.

44. Riedl J, Flynn KC, Raducanu A, Gartner F, Beck G, Bosl M, et al. Lifeact mice for studying F-actin dynamics. Nat Methods. 2010;7(3):168–9.

45. Muzumdar MD, Tasic B, Miyamichi K, Li L, and Luo L. A global double-fluorescent Cre reporter mouse. Genesis. 2007;45(9):593–605.

46. Zhang Y, Conti MA, Malide D, Dong F, Wang A, Shmist YA, et al. Mouse models of MYH9-related disease: mutations in nonmuscle myosin II-A. Blood. 2012;119(1):238–50.

47. Franco CA, Jones ML, Bernabeu MO, Vion AC, Barbacena P, Fan J, et al. Non-canonical Wnt signalling modulates the endothelial shear stress flow sensor in vascular remodelling. Elife. 2016;5:e07727.

48. Burbelo PD, Drechsel D, and Hall A. A conserved binding motif defines numerous candidate target proteins for both Cdc42 and Rac GTPases. J Biol Chem. 1995;270(49):29071–4.

